# EukaUTR: a foundation model unifying functional modelling and design of eukaryotic 3^′^ UTRs

**DOI:** 10.64898/2026.09.07.749809

**Authors:** Mei Lang, Xingyu Fang, Mingxuan Chen, Zhen Wang, Zhaowen Cheng, Xiagu Zhu, Kin Yip Tam, Junwei Zhang, Xiaolin Li

## Abstract

Eukaryotic mRNA 3^′^ UTRs encode regulatory information that shapes post-transcriptional control, RNA fate and gene expression. However, a 3^′^ UTR-specific foundation model that spans broad eukaryotic sequence diversity while supporting both functional prediction and sequence design is lacking. Here we present EukaUTR, a 3^′^ UTR-specific foundation model trained on a large-scale eukaryotic 3^′^ UTR sequence corpus spanning diverse evolutionary lineages. Across 13 prediction tasks spanning post-transcriptional regulation, RNA fate and expression output, EukaUTR models matched or exceeded the strongest external baselines on nearly all tasks, with relative improvements of up to 27.45%. EukaUTR also generated de novo 3^′^ UTRs with natural-like sequence and regulatory properties. EukaUTR-Guide further derived stability-associated signals from small sequence sets to guide editing towards enhanced predicted stability. Together, EukaUTR provides a sequence-to-function-to-design framework for transferable 3^′^ UTR modelling and design.

## 1 Introduction

The 3^′^ untranslated region (3′ UTR) is a major cis-regulatory region of eukaryotic mRNAs that shapes post-transcriptional gene regulation. Through interactions with RNA-binding proteins, microRNAs and other regulatory factors, 3′ UTRs influence mRNA stability, localization, translation and protein output, whereas alternative polyadenylation alters this regulatory potential by changing 3′ UTR length and cis-regulatory element content [1–6]. Although 3′ UTRs contain diverse regulatory elements, how these elements collectively shape RNA fate remains poorly understood. Deciphering how regulatory information is encoded in 3′ UTR sequence is therefore important for understanding post-transcriptional regulation, interpreting non-coding variation and designing mRNAs with desired properties.

Experimental studies have revealed multiple mechanisms of 3′ UTR regulation, including RNA-binding protein interactions, RNA modifications, microRNA targeting and alternative polyadenylation, as well as sequence features that influence mRNA stability and translation [1, 2, 4–8]. However, most experimental approaches examine specific regulatory factors, sequence elements or cell line, limiting systematic exploration of the broader 3′ UTR regulatory landscape [2, 9, 10]. Computational models have broadened these efforts by leveraging predefined sequence features, predicted RNA structural features and assay-specific datasets, but many remain tailored to individual regulatory processes or experimental settings, limiting their ability to learn transferable representations from 3′ UTR sequence [9, 11, 12]. This limitation is particularly important for 3′ UTRs, where the regulatory effect of a cis-regulatory element is shaped by its surrounding sequence and structural context rather than by the element in isolation [4]. This context dependence motivates models that can learn regulatory information directly from 3′ UTR sequence and transfer it across diverse functional tasks.

Self-supervised nucleotide language models have shown that large-scale sequence pre-training can learn transferable representations that support diverse DNA-and RNA-sequence prediction tasks [13–16]. Building on this progress, 3′ UTR-specific language models have shown that pretrained sequence representations can capture regulatory information from 3′ UTR sequence. For example, 3UTRBERT [17], pretrained on human 3′ UTR sequences, learned interpretable representations that supported prediction across diverse 3′ UTR regulatory tasks, whereas GEMORNA [18], trained on mammalian 3′ UTRs, extended UTR modelling to de novo 3′ UTR generation. In parallel, mRNABERT [19], pretrained on full-length mRNA sequences, learned integrated transcript-level representations that supported both region-specific and full-length mRNA prediction tasks, including several 3′ UTR-related tasks. However, existing 3′ UTR-focused language models remain largely restricted to human or mammalian sequence space. Although recent models have extended UTR modelling from functional prediction to sequence generation, a 3′ UTR-specific framework that combines broad eukaryotic representation learning with transferable functional prediction and sequence design is still lacking. These limitations motivate a 3′ UTR-specific foundation model pretrained across broad eukaryotic sequence diversity and capable of supporting transferable functional prediction and sequence design.

Here we introduce EukaUTR, a 3′ UTR-specific foundation model pretrained across broad eukaryotic sequence diversity. Pre-training first used approximately 7.0 million non-redundant 3′ UTRs from diverse eukaryotic lineages, followed by continued pre-training on a high-confidence vertebrate corpus comprising species excluded from the first stage. This two-stage design couples broad eukaryotic representation learning with subsequent refinement on high-confidence vertebrate 3′ UTRs. We evaluated EukaUTR across 13 downstream prediction tasks covering post-transcriptional regulation, RNA stability and decay, and expression output. We then tested whether the model captured local cis-regulatory effects and sequence regions that are sensitive to perturbation. Beyond functional prediction and regulatory analysis, we assessed EukaUTR for de novo 3′ UTR generation and characterized the resulting sequences across sequence, structural and regulatory properties. Finally, we developed EukaUTR-Guide, which derives stability-associated signals from small sets of 3′ UTRs with different stability properties and uses them to guide sequence editing towards improved predicted stability. Together, these results position EukaUTR as a unified sequence-to-function-to-design framework that connects broad eukaryotic 3′ UTR representation learning with functional prediction, regulatory analysis and sequence design. These capabilities may enable more systematic, model-guided design of 3′ UTRs for mRNA therapeutics.

## 2 Results

### 2.1 EukaUTR learns representations across broad eukaryotic 3**′** UTR sequence space

We developed EukaUTR, a foundation model for eukaryotic 3′ UTRs. EukaUTR supports functional prediction, local cis-regulatory analysis, de novo 3′ UTR generation and property-directed sequence editing (Fig. 1a). Pre-training proceeded in two stages using annotated 3′ UTRs from Ensembl and Ensembl Genomes [20, 21]. Both corpora underwent the same sequence-length filtering and MMseqs2-based redundancy reduction at 80% sequence identity [22].

**Figure 1:**
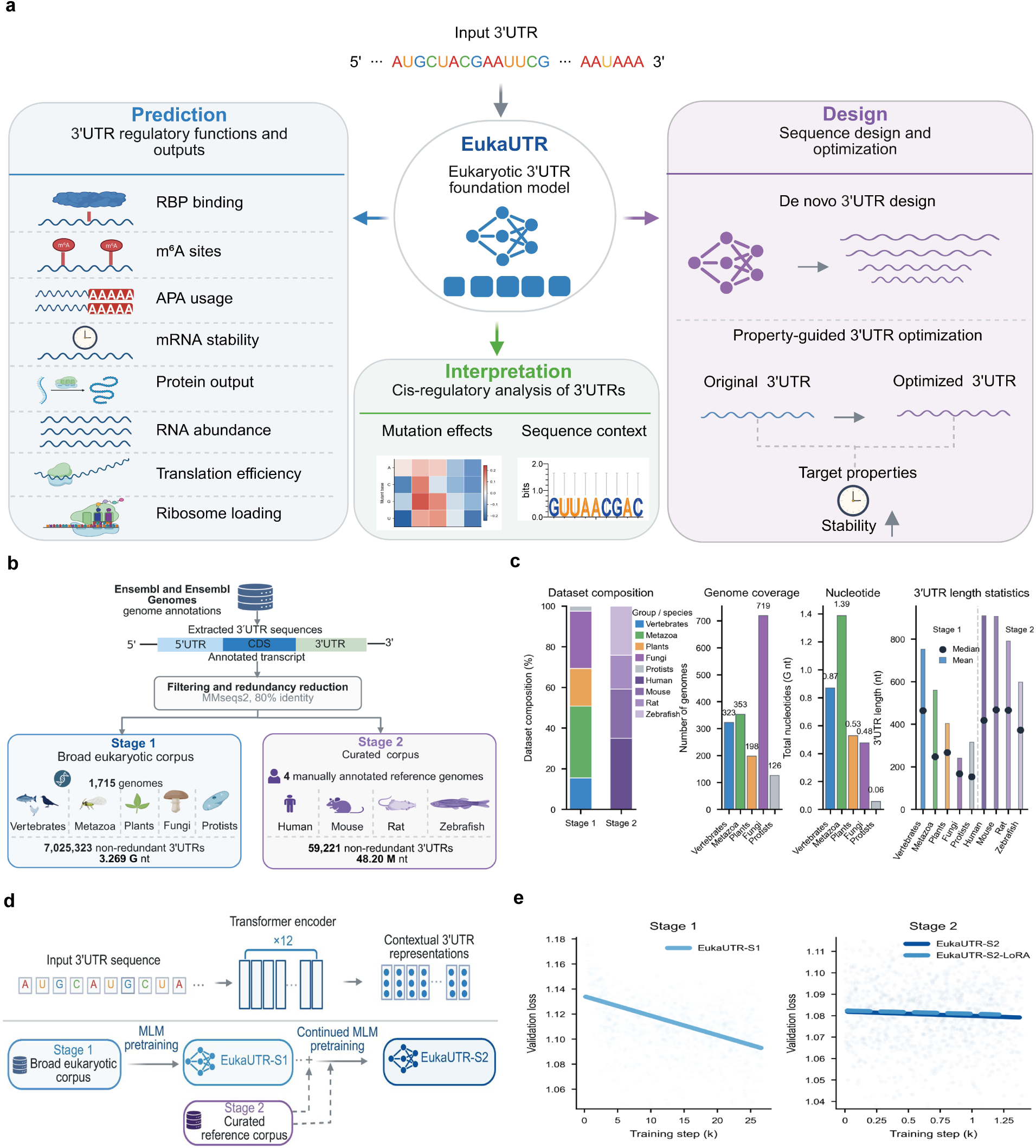
Overview of EukaUTR. **a,** Applications of EukaUTR to functional prediction, local cis-regulatory analysis, de novo 3′ UTR generation and property-directed sequence editing. **b,** Pre-training corpora, with Stage 1 spanning broad eukaryotic diversity and Stage 2 focusing on four vertebrate species excluded from Stage 1. **c,** Summary of the Stage 1 and Stage 2 corpora, including taxonomic composition, number of genomes, total nucleotide content and 3′ UTR length distributions. **d,** Two-stage pre-training of EukaUTR, with Stage 1 learning from broad eukaryotic 3′ UTR diversity and Stage 2 refining the model on high-confidence vertebrate sequences using either full-parameter continued pre-training or low-rank adaptation. **e,** Validation-loss trends during Stage 1 pre-training and the two Stage 2 training strategies

Stage 1 provided broad eukaryotic sequence diversity for representation learning, comprising approximately 7.03 million non-redundant 3′ UTRs from 1,715 genomes and 3.27 billion nucleotides. Stage 2 comprised 59,221 high-confidence 3′ UTRs from human, mouse, rat and zebrafish, all of which were excluded from Stage 1 (Fig. 1b,c; Supplementary Figs. 1, 2; Supplementary Tables 1, 2). EukaUTR uses a 12-layer Transformer encoder [23–25] trained with a masked language-modelling objective to learn contextual representations of 3′ UTR sequences (Fig. 1d). Stage 1 pre-training produced EukaUTR-S1, which was subsequently further trained on the Stage 2 corpus of high-confidence 3′ UTRs through either full-parameter continued pre-training or low-rank adaptation [26], yielding EukaUTR-S2 and EukaUTR-S2-LoRA, respectively. Validation loss decreased during Stage 1 pre-training, and both Stage 2 refinement strategies produced further reductions in validation loss and perplexity on the Stage 2 validation set (Fig. 1e; Supplementary Fig. 3). We subsequently evaluated all three models across regulatory and RNA-fate prediction tasks, as well as de novo 3′ UTR generation.

### 2.2 EukaUTR predicts diverse modes of post-transcriptional regulation

#### EukaUTR predicts RNA-binding protein interactions

RNA-binding proteins bind specific sites within 3′ UTRs, with binding specificity shaped by local sequence and RNA structure, thereby contributing to post-transcriptional regulation [4, 27]. To evaluate RBP-binding prediction, we benchmarked EukaUTR-S1, EukaUTR-S2 and EukaUTR-S2-LoRA against a broad panel of existing methods, including task-specific supervised models, general RNA and nucleotide language models, and 3′ UTR-specific language models (Fig. 2a,b). We evaluated RBP-binding prediction in two settings: a fivefold cross-validated eCLIP benchmark spanning 22 RBP–cell-line combinations and a gene-held-out CLIP benchmark comprising 31 experiments, in which test sequences originated from genes excluded from downstream model fitting [17].

**Figure 2:**
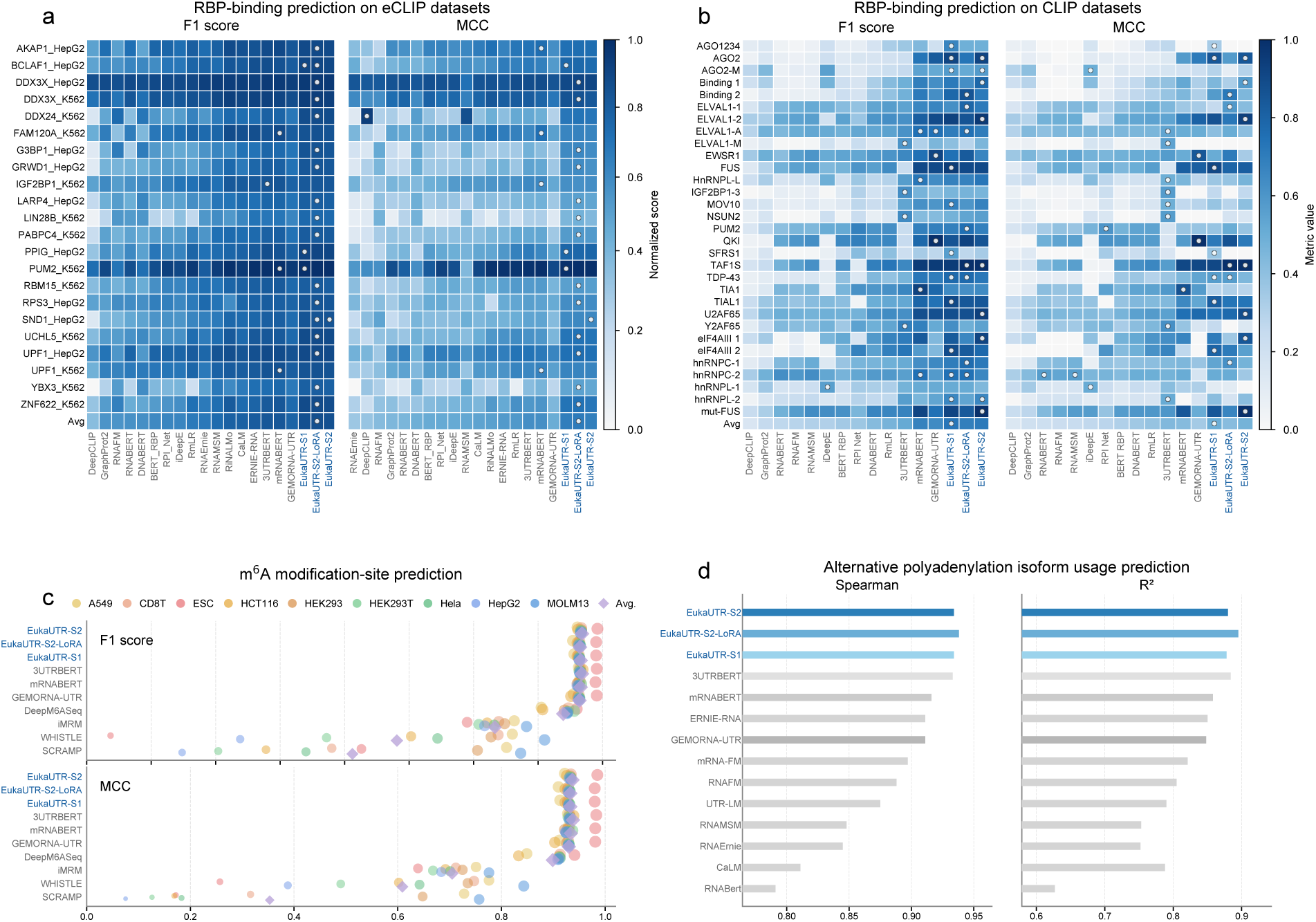
EukaUTR predicts diverse modes of post-transcriptional regulation. **a,b,** RBP-binding prediction on the eCLIP **(a)** and gene-held-out CLIP **(b)** benchmarks. Heatmaps show F1 scores and Matthews correlation coefficients (MCCs) across individual RBP-binding tasks, with the bottom row showing macro-averaged performance. White markers indicate the highest-performing model for each task. **c,** m^6^A-site prediction across nine human cell lines. Circles represent individual cell lines and diamonds represent the macro-average. **d,** Prediction of proximal polyadenylation-site usage.

On the eCLIP benchmark, all three EukaUTR models outperformed the strongest external baselines in macro-averaged F1 and MCC across the 22 RBP tasks. EukaUTR-S2-LoRA achieved the highest macro-averaged F1 and MCC, reaching 0.782 and 0.567, respectively, while EukaUTR-S1 and EukaUTR-S2 showed comparable performance. Across the three EukaUTR models, relative improvements over the strongest external baselines ranged from 3.46% to 4.13% for F1 and from 10.74% to 12.72% for MCC (Fig. 2a; Supplementary Tables 4 and 5). At the individual-task level, EukaUTR-S2-LoRA ranked first or jointly first in 17 of 22 tasks for F1 and 13 of 22 for MCC, whereas EukaUTR-S1 did so in three tasks for each metric and EukaUTR-S2 in one task for each metric.

On the gene-held-out CLIP benchmark, which comprised 31 experiments spanning 19 RBPs, EukaUTR-S1 achieved the highest macro-averaged performance, with an F1 score of 0.723 and an MCC of 0.497 (Fig. 2b; Supplementary Tables 6 and 7). All three EukaUTR models achieved higher F1 scores than the strongest external baseline, with relative improvements ranging from 1.78% to 6.95%. For MCC, EukaUTR-S2 and EukaUTR-S1 achieved improvements of 7.31% and 13.47%, respectively, whereas EukaUTR-S2-LoRA showed a lower macro-averaged MCC than the strongest external baseline. At the individual-experiment level, EukaUTR-S1 ranked first or jointly first in 11 of 31 experiments for F1 and 7 of 31 for MCC.

Together, these results show strong RBP-binding prediction across all three EukaUTR models, with the benefit of Stage 2 adaptation depending on the evaluation setting. EukaUTR-S2-LoRA performed best on eCLIP, whereas EukaUTR-S1 performed best on the more stringent gene-held-out CLIP benchmark.

#### EukaUTR predicts m^6^A sites

N^6^-methyladenosine (m^6^A) is a widespread modification of eukaryotic mRNA that influences RNA processing, stability and translation [7, 8]. To evaluate whether EukaUTR could identify m^6^A-modified adenosines from sequence, we benchmarked EukaUTR-S1, EukaUTR-S2 and EukaUTR-S2-LoRA against representative models, including task-specific m^6^A predictors, general mRNA language models and 3′ UTR-specific language models (Fig. 2c). The benchmark covered nine human cell lines, with each evaluated using ten approximately balanced positive–negative subsets following the 3UTRBERT protocol [17] (Supplementary Tables 8 and 9).

Across the nine cell lines, all three EukaUTR models showed strong and similar overall performance. EukaUTR-S2 achieved the highest macro-averaged scores among the three models, with an F1 score of 0.968 and an MCC of 0.937, tying 3UTRBERT for the highest F1 and closely approaching its MCC of 0.938. Across individual contexts, EukaUTR-S2 ranked first in six of nine for F1 and five of nine for MCC, and ranked among the top two models in seven contexts for both metrics. EukaUTR-S1 and EukaUTR-S2-LoRA showed comparable macro-averaged performance, with F1 scores of 0.966 and 0.965, respectively, and an MCC of 0.933 for both models. Together, these results show that m^6^A-site prediction remained consistently strong across all three EukaUTR models, with only small differences in overall performance.

#### EukaUTR predicts proximal polyadenylation site usage

Alternative polyadenylation (APA) generates mRNA isoforms with distinct 3′ ends, altering 3′ UTR length and regulatory-element composition and thereby influencing mRNA stability and translation, with consequences for protein output [5, 6, 28]. We therefore evaluated whether EukaUTR could predict proximal polyadenylation-site usage using the BEACON APA benchmark, derived from the APARENT massively parallel reporter assay [29, 30] (Fig. 2d; Supplementary Table 10). All three EukaUTR models showed similar performance. EukaUTR-S2-LoRA achieved the highest *R*^2^ and Spearman correlation, reaching 0.895 and 0.938, respectively, corresponding to relative gains of 1.24% and 0.54% over the strongest external baseline, 3UTRBERT. EukaUTR-S2 and EukaUTR-S1 showed closely similar performance, with Spearman correlations of 0.934 for both models and *R*^2^ values of 0.880 and 0.878, respectively.

### 2.3 EukaUTR predicts mRNA stability and gene-expression outputs

3′ UTR sequence encodes regulatory information that influences mRNA stability, abundance and translation, thereby shaping quantitative gene-expression outputs [2, 3, 9, 31]. We therefore evaluated EukaUTR for quantitative prediction of mRNA stability, abundance, translation and protein output (Fig. 3a–d). We therefore compared EukaUTR with 3UTRBERT [17], GEMORNA-UTR [18] and mRNABERT [19], three language models applicable to 3′ UTR modelling that showed strong performance in the preceding post-transcriptional regulation benchmarks. We evaluated EukaUTR across mRNA stability, RNA abundance, translational efficiency and protein output in diverse reporter and cell lines [2, 3, 9, 32, 33].

**Figure 3:**
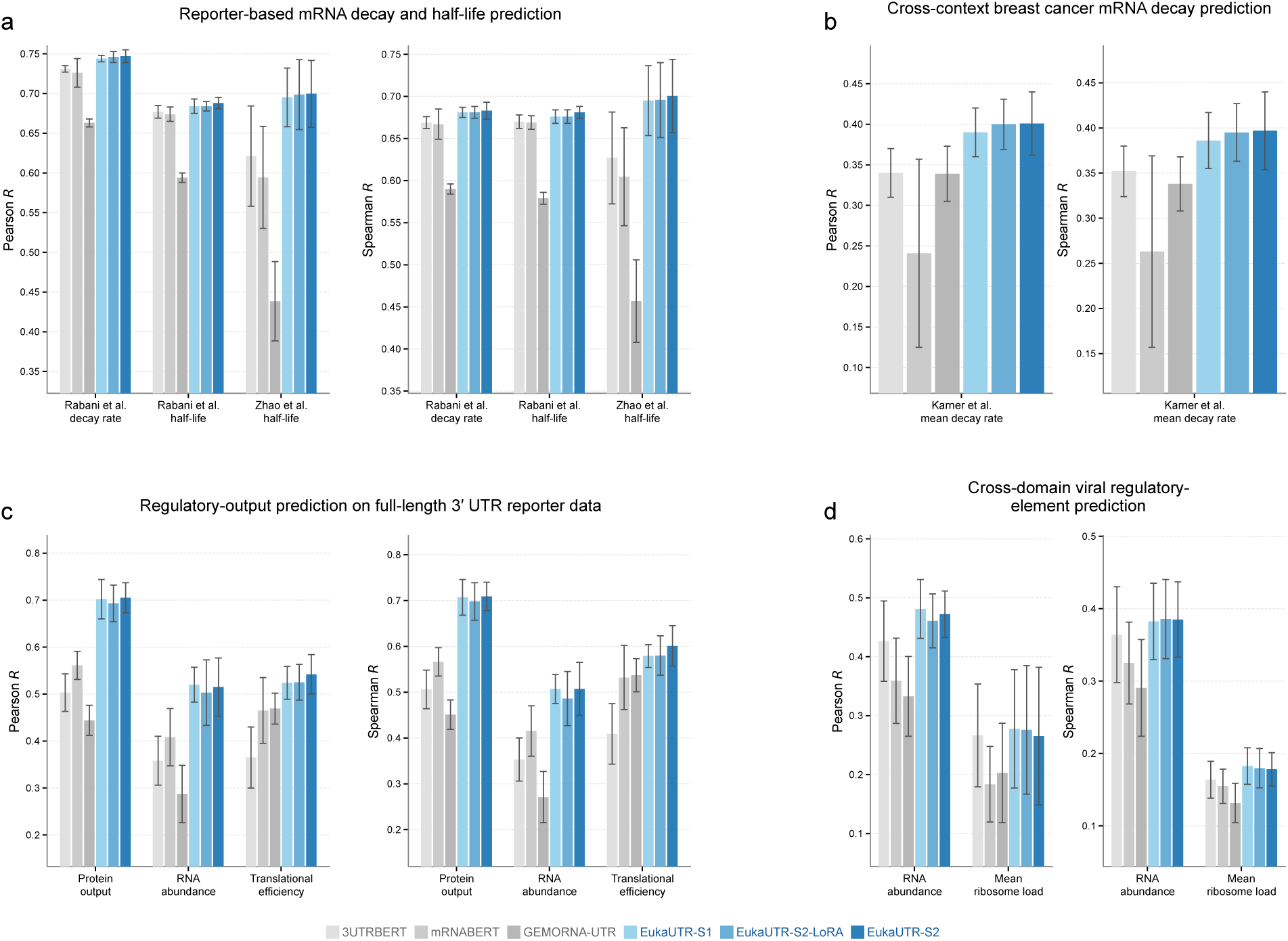
EukaUTR predicts mRNA stability and expression outputs. **a,** Prediction of reporter mRNA degradation rate and half-life across three MPRA-derived benchmarks from Rabani et al. and Zhao et al. **b,** Prediction of an mRNA decay phenotype averaged across six breast cancer cell lines using data from Karner et al. **c,** Prediction of protein output, RNA abundance and translational efficiency using a full-length human 3′ UTR MPRA dataset. Performance values are averaged across the CAG and PGK promoter contexts. **d,** Prediction of RNA abundance and mean ribosome load (MRL) for heterologous viral sequences assayed in a common 3′ UTR reporter context. Bars show mean Pearson and Spearman correlations; error bars denote s.d. across folds.

#### EukaUTR predicts mRNA stability across diverse experimental systems

To assess whether EukaUTR captures sequence determinants of mRNA stability, we evaluated it across complementary mRNA-stability benchmarks derived from reporter assays and cellular transcriptomic data (Fig. 3a,b). Reporter-based evaluation included zebrafish mRNA degradation rate and half-life from Rabani et al. [2], and human reporter mRNA half-life from Zhao et al. [9]. We further evaluated an mRNA decay phenotype using data from Karner et al. [32].

On the Rabani et al. UTR-seq benchmarks, all three EukaUTR models showed strong predictive performance. EukaUTR-S2 achieved the highest correlations for both targets, reaching *r* = 0.747 and *ρ* = 0.683 for degradation-rate prediction and *r* = 0.688 and *ρ* = 0.681 for derived half-life prediction. Relative to the strongest external baseline, 3UTRBERT, these values corresponded to gains of 2.19% and 2.09% in Pearson and Spearman correlation for degradation rate, and 1.62% and 1.64% for half-life, respectively (Supplementary Tables 11 and 12). EukaUTR-S1 and EukaUTR-S2-LoRA also outperformed 3UTRBERT for both targets and both correlation metrics, with these improvements supported by paired fold-level comparisons (two-sided paired Wilcoxon signed-rank tests, Holm-adjusted *P* ≤ 0.0059).

On the Zhao et al. fast-UTR half-life benchmark, EukaUTR-S2 achieved the highest correlations, reaching *r* = 0.700 and *ρ* = 0.700, corresponding to relative gains of 12.72% and 11.64%, respectively, over the strongest external baseline, 3UTRBERT (Supplementary Table 13). EukaUTR-S1 and EukaUTR-S2-LoRA showed closely comparable performance (*r* = 0.695–0.699; *ρ* = 0.695–0.696). All three EukaUTR models outperformed the external baselines for both correlation metrics in paired fold-level comparisons (two-sided paired Wilcoxon signed-rank tests, Holm-adjusted *P* = 0.0176 for all comparisons), whereas no significant differences were detected among the three EukaUTR models (*P*_adj_ = 1.0).

On the Karner et al. mRNA decay benchmark, all three EukaUTR models showed strong predictive performance. EukaUTR-S2 achieved the highest mean correlations, reaching *r* = 0.401 and *ρ* = 0.397, corresponding to relative gains of 17.94% and 12.78%, respectively, over the strongest external baselines (Fig. 3b; Supplementary Table 14). EukaUTR-S1 and EukaUTR-S2-LoRA showed closely comparable performance. In paired fold-level comparisons, all three EukaUTR models significantly outperformed the strongest external baseline in Pearson correlation, whereas a significant improvement in Spearman correlation was detected only for EukaUTR-S2-LoRA.

Together, these results show that EukaUTR supports quantitative prediction of mRNA stability across complementary reporter-based and mRNA decay benchmarks, with consistently strong performance across all three EukaUTR models.

#### EukaUTR predicts expression outputs across diverse reporter systems

To assess whether EukaUTR could predict expression outputs beyond mRNA stability, we evaluated the full-length human 3′ UTR MPRA of West et al. [3], which measured protein output, RNA abundance and translational efficiency under CAG and PGK promoter contexts; performance was averaged across the two promoter contexts (Fig. 3c; Supplementary Table 15). Across the three outputs, EukaUTR models consistently outperformed the external baselines. For protein output, EukaUTR-S2 achieved the highest correlations (*r* = 0.705, *ρ* = 0.709), corresponding to relative gains of 25.67% and 25.27%, respectively, over the strongest external baseline, mRNABERT [19]. For RNA abundance, EukaUTR-S1 achieved the highest Pearson correlation (*r* = 0.520), whereas EukaUTR-S1 and EukaUTR-S2 jointly achieved the highest Spearman correlation (*ρ* = 0.507), representing relative gains of 27.45% and 22.17%, respectively, over mRNABERT. For translational efficiency, EukaUTR-S2 achieved the highest correlations (*r* = 0.542, *ρ* = 0.601), with relative gains of 15.57% and 11.92% over the strongest external baseline, GEMORNA-UTR. Across all three outputs, each EukaUTR model significantly outperformed the corresponding strongest external baseline for both Pearson and Spearman correlations (two-sided paired Wilcoxon signed-rank tests, Holm-adjusted *P* ≤ 0.0195).

To assess whether EukaUTR could generalize beyond eukaryotic 3′ UTR sequences, we evaluated the viromics MPRA of Seo et al. [33], which measured RNA abundance and mean ribosome load (MRL) for viral sequences in a reporter assay (Fig. 3d; Supplementary Table 16). For RNA abundance, EukaUTR-S1 achieved the highest mean Pearson correlation (*r* = 0.481), whereas EukaUTR-S2-LoRA achieved the highest mean Spearman correlation (*ρ* = 0.386), corresponding to relative gains of 12.91% and 6.04%, respectively, over the strongest external baseline, 3UTRBERT. Both improvements remained significant after Holm correction (*P*_adj_ = 0.0176 and 0.0293, respectively). For MRL, EukaUTR-S1 achieved the highest mean correlations (*r* = 0.278, *ρ* = 0.183), corresponding to relative gains of 4.12% and 11.59%, respectively, over 3UTRBERT, with the Spearman but not the Pearson difference remaining significant after Holm correction (*P*_adj_ = 0.0176 for Spearman).

Across the nine mRNA-stability and expression-output tasks, EukaUTR models consistently outperformed the strongest external baselines, with significant improvements in both Pearson and Spearman correlations for eight of the nine tasks. EukaUTR-S2 achieved the highest mean correlations in six tasks, including all four mRNA-stability benchmarks, while EukaUTR-S2-LoRA matched or exceeded EukaUTR-S1 on several tasks. Together, these results indicate that high-confidence Stage 2 refinement was particularly beneficial for mRNA-stability prediction, although its effect varied across downstream tasks.

### 2.4 EukaUTR captures local cis-regulatory effects and regulatory sequence context

To test whether EukaUTR resolves local regulatory information underlying its transcript-level predictions, we examined two complementary settings (Fig. 4). Because EukaUTR-S2 achieved the highest mean correlations across the preceding mRNA-stability benchmarks, we used this model for the subsequent local cis-regulatory analyses. A half-life predictor fine-tuned on the Zhao et al. human reporter benchmark was applied to a separate *CXCL2* saturation-mutagenesis dataset to assess local mutational effects, while a degradation-rate predictor fine-tuned on the Rabani et al. zebrafish UTR-seq benchmark was applied to RBMS3-bound reporter elements and endogenous CLIP-site windows to examine RBMS3-associated sequence context.

**Figure 4:**
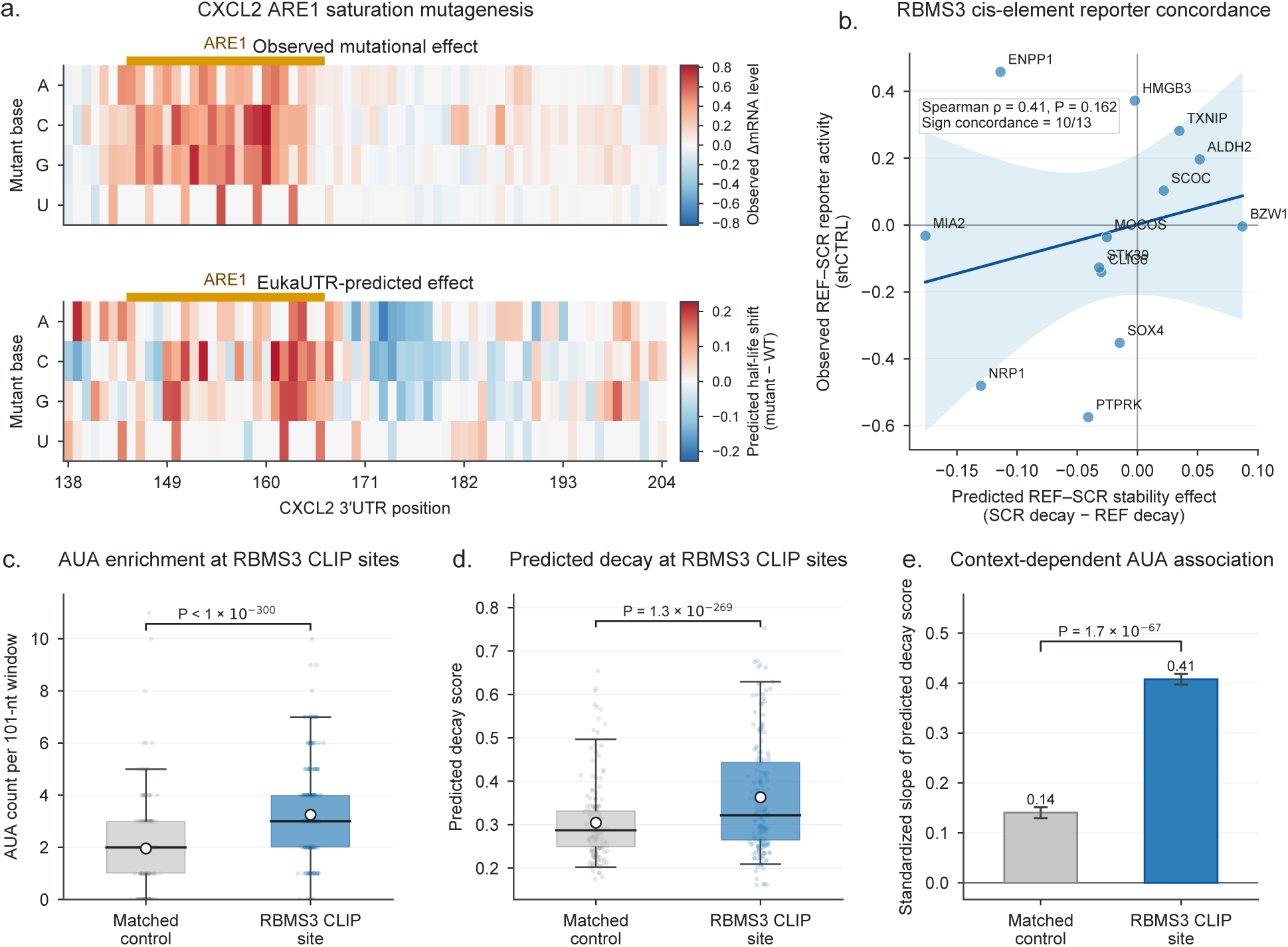
EukaUTR captures local cis-regulatory effects and sequence context. **a,** Saturation mutagenesis of 67-nt region of the *CXCL2 3′* UTR containing the 21-nt ARE1 element [9]. Heat maps show experimentally measured changes in reporter mRNA abundance and EukaUTR-S2-predicted changes in mRNA half-life relative to wild type across all 201 possible single-nucleotide substitutions. **b,** Comparison of EukaUTR-S2-predicted stability effects and experimentally measured reporter effects for 13 RBMS3-bound reference elements and their dinucleotide-preserving scrambled controls [32]. Line and shading indicate the linear fit and 95% confidence interval. c,d, AUA counts **(c)** and EukaUTR-S2-predicted decay scores **(d)** in RBMS3 CLIP-site windows compared with matched control windows from the same 3′ UTRs (n = 11,248 pairs). **e,** Association between AUA density and EukaUTR-S2-predicted decay scores in RBMS3-bound and matched-control windows. Error bars denote cluster-robust standard errors.

We first analysed all 201 possible single-nucleotide substitutions across a 67-nt region of the *CXCL2* 3′ UTR comprising the 21-nt ARE1 element and 46 nt of flanking sequence [9]. Experimental effects were defined as changes in steady-state reporter mRNA abundance relative to the wild-type sequence, whereas model effects were defined as changes in EukaUTR-S2-predicted mRNA half-life relative to wild type (Fig. 4a). Across these substitutions, predicted and experimentally measured effects showed modest but significant correlations (Pearson’s *r* = 0.210, *P* = 2.83 × 10^−3^; Spearman’s *ρ* = 0.219, *P* = 1.80 × 10^−3^). Predicted effect magnitudes were also greater within ARE1 than in the flanking sequence (mean absolute effect, 0.0817 versus 0.0572; two-sided Wilcoxon rank-sum test, *P* = 0.024), indicating that EukaUTR-S2 localized elevated mutational sensitivity to the ARE1 region. Although EukaUTR-S2 localized elevated mutational sensitivity to ARE1, it did not reliably resolve the effects of individual substitutions within the element.

Karner et al. identified RBMS3 as a stabilizing regulator of AUA-containing transcripts and showed that RBMS3-bound regions in 3′ UTRs are enriched for AUA-rich sequence elements [32]. We therefore asked whether EukaUTR-S2, fine-tuned for mRNA degradation-rate prediction, captured the corresponding cis-sequence signature. In a dual-reporter assay of 13 RBMS3-bound reference elements and their dinucleotide-preserving scrambled controls, EukaUTR-S2-predicted stability effects were positively, but not significantly, correlated with experimentally measured reporter effects (Spearman’s *ρ* = 0.412, *P* = 0.162; Pearson’s *r* = 0.232, *P* = 0.446; Fig. 4b). Predicted and experimental effect directions agreed in 10 of the 13 element pairs. These results suggest directional concordance, but the small sample size and non-significant correlations do not support reliable quantitative prediction of effect magnitude.

To test the underlying sequence signal at larger scale, we next analysed 11,248 pairs of 101-nt windows centred on endogenous RBMS3 CLIP sites and matched control regions from the same 3′ UTRs. Consistent with the reported AUA-rich sequence preference of RBMS3, bound windows contained significantly higher AUA counts than matched controls (two-sided paired Wilcoxon signed-rank test, *P* < 1 × 10^−300^; Fig. 4c). RBMS3-bound windows also showed higher EukaUTR-S2-predicted decay scores (*P* = 1.3 × 10^−269^; Fig. 4d), indicating that the model responds systematically to sequence features enriched at RBMS3-binding sites.

We then asked whether this model response was related to AUA enrichment itself. The association between AUA density and predicted decay was substantially stronger in RBMS3-bound windows than in matched controls, with standardized slopes of 0.41 and 0.14, respectively. This difference was supported by a significant interaction between AUA density and window class (*P* = 1.7 × 10^−67^; Fig. 4e). Thus, EukaUTR-S2 captured an AUA-rich sequence signal associated with its decay predictions at RBMS3-bound regions.

Because EukaUTR receives sequence alone and does not model RBMS3 abundance, binding occupancy or cellular state, the higher predicted decay scores at RBMS3-bound regions should not be interpreted as evidence that RBMS3 promotes mRNA degradation. Rather, they indicate that EukaUTR-S2 captures a sequence-encoded signal associated with decay propensity, whereas the realized regulatory effect in cells depends on the trans-acting activity of RBMS3 and the broader cellular context.

Together, these analyses show that EukaUTR-S2 captures regional and context-dependent cis-regulatory sequence signals beyond transcript-level prediction, while supporting regional rather than nucleotide-level or mechanistic interpretation.

### 2.5 EukaUTR generates de novo 3**′** UTRs with natural-like sequence properties

We next asked whether EukaUTR could generate de novo 3′ UTRs with properties resembling those of natural 3′ UTRs. Given the multiple interrelated properties of natural 3′ UTRs, including nucleotide composition, local sequence organization, structural propensity and cis-regulatory content, we evaluated the generated sequences across complementary sequence-level criteria. We first sampled natural 3′ UTRs from the full pre-training corpus (Natural), whose lengths defined a common set of target lengths spanning 100–1,022 nt. The same sampled target lengths were used to generate equiprobable random nucleotide sequences (Random) and sequences from EukaUTR-S1, EukaUTR-S2, EukaUTR-S2-LoRA, 3UTRBERT [17], mRNABERT [19] and GEMORNA-UTR [18] (Fig. 5).

**Figure 5:**
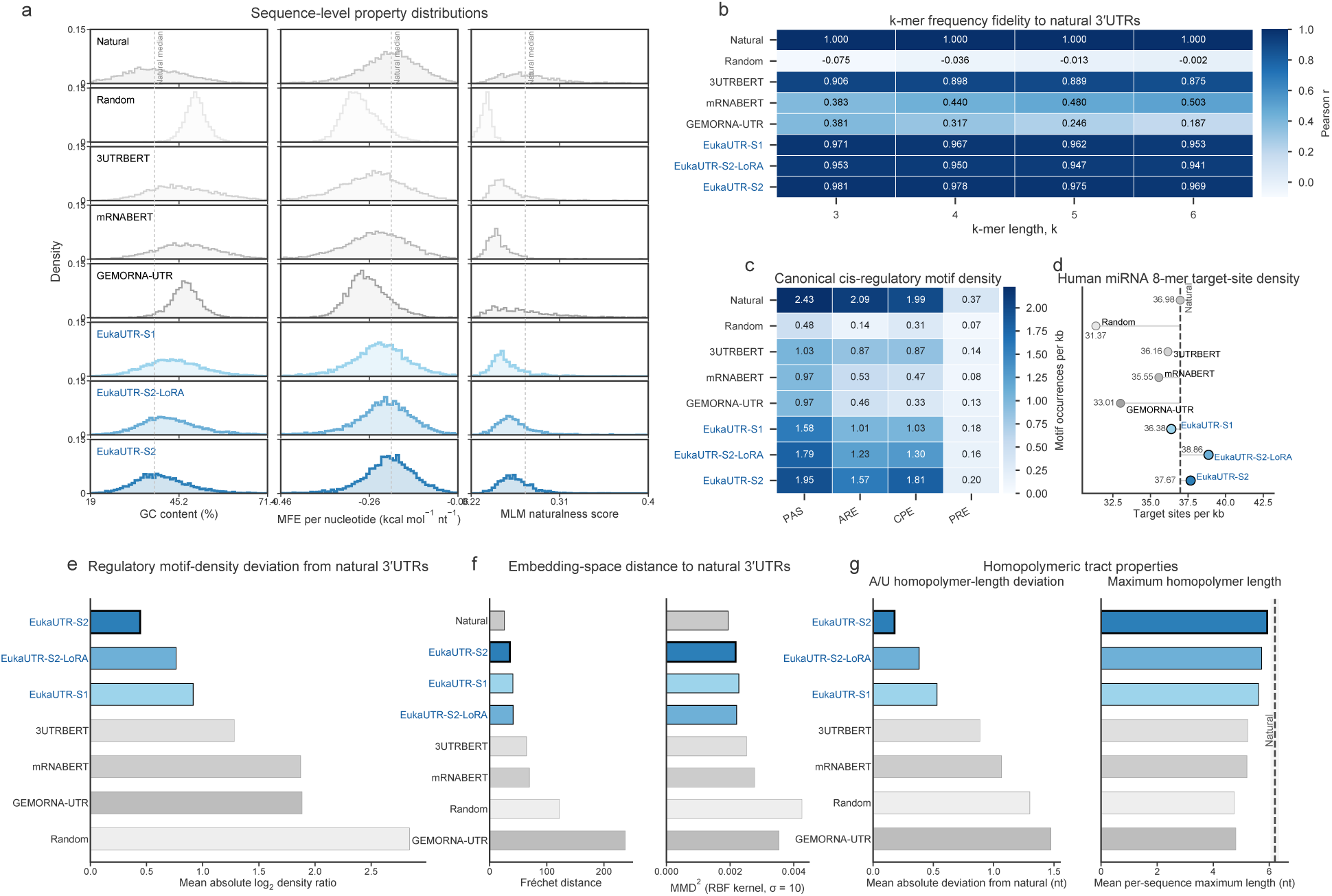
EukaUTR generates de novo 3′ UTRs with natural-like sequence properties. Natural sequences, generated sequences and random controls were length matched. **a,** Distributions of GC content, length-normalized RNAfold-predicted minimum free energy (MFE) and EukaUTR-S2 naturalness scores. **b,** Pearson correlations between generated and natural *k-*mer frequency profiles for *k* = 3–6. **c,** Densities of canonical polyadenylation signals (PAS), AU-rich elements (AREs), cytoplasmic polyadenylation elements (CPEs) and Pumilio response elements (PREs). **d,** Density of predicted human miRNA 8-mer target sites. **e,** Mean absolute log2 deviation of canonical motif densities from natural sequences. **f,** Fréchet distance and squared maximum mean discrepancy (MMD2) between generated and natural sequences in EukaUTR-S2 embedding space. **g**, A/U homopolymer properties of generated and natural sequences.

All three EukaUTR models generated sequences with natural-like GC content, length-normalized minimum free energy and EukaUTR-S2-based naturalness scores (Fig. 5a). They also closely matched natural local sequence composition, with 3–6-mer frequency correlations of *r* = 0.941–0.981 across the three models. EukaUTR-S2 achieved the highest correlations at each *k* (*r* = 0.969–0.981), exceeding the strongest external comparator, 3UTRBERT (*r* = 0.875–0.906) (Fig. 5b). This similarity extended to regulatory sequence content: across canonical PAS, ARE, CPE and PRE motifs, all three EukaUTR-generated sets more closely approximated natural motif densities than the external comparators, with EukaUTR-S2 showing the smallest overall deviation, followed by EukaUTR-S2-LoRA and EukaUTR-S1 (Fig. 5c,e). Predicted human miRNA 8-mer target-site densities were likewise close to the natural reference across all three models (Fig. 5d). Although canonical motif densities generally remained below natural levels, these results show that EukaUTR-generated sequences captured multiple sequence-level properties of natural 3′ UTRs.

We next asked whether this similarity was also reflected in representation space. In EukaUTR-S2 embedding space, sequences generated by all three EukaUTR models were closer to the length-matched natural reference than those generated by the external comparators according to both Fréchet distance and MMD^2^. EukaUTR-S2 showed the smallest distances, while EukaUTR-S1 and EukaUTR-S2-LoRA also remained close to the natural reference (Fig. 5f). As these metrics were computed in EukaUTR-S2 embedding space, they were interpreted as internal measures of representation-space proximity and considered alongside the direct sequence-level analyses.

Finally, we examined whether the generated sequences showed evidence of low-complexity degeneration. Across the three EukaUTR models, A-and U-homopolymer lengths and maximum per-sequence homopolymer lengths remained close to those of natural 3′ UTRs (Fig. 5g). The fractions of sequences exceeding the natural-reference 95th percentile for homopolymer length or containing homopolymeric tracts of ≥6 nt were likewise low (Supplementary Fig. 4a,b).

We then assessed whether the generated sequences showed high-coverage redundancy within each generated set or similarity to the EukaUTR pre-training corpus using MMseqs2 [22]. At ≥80% sequence identity and ≥0.8 alignment coverage, no qualifying non-self matches were detected within any of the three EukaUTR-generated sets. At the same thresholds, a small number of qualifying within-set matches were detected for GEMORNA-UTR [18] (Supplementary Fig. 4c). No qualifying matches to the complete EukaUTR pre-training corpus were detected for sequences generated by any method.

Together, these analyses show that all three EukaUTR models generated de novo 3′ UTRs with multiple natural-like sequence properties, without evidence of substantial low-complexity degeneration, high-coverage within-set redundancy or high-coverage similarity to the pre-training corpus. These computational properties do not, however, establish biological function.

### 2.6 EukaUTR-Guide enables property-directed 3**′** UTR editing

To enable precise design of 3′ UTRs with desired properties, we developed EukaUTR-Guide, which derives a property-associated signal from sequences with contrasting predicted properties and uses this signal to guide targeted sequence editing (Fig. 6a). We then used EukaUTR-Guide for predicted mRNA-stability optimization by deriving a stability-associated signal from 100 3′ UTR fragments with low predicted degradation rates and 100 with high predicted degradation rates. This signal was used to prioritize nucleotide positions that deviated most from the low-degradation profile. These positions were then masked, and EukaUTR-S2 predicted context-compatible replacement nucleotides. EukaUTR-Guide therefore uses the property-associated signal to prioritize editing sites, while EukaUTR-S2 predicts context-compatible replacement nucleotides. No additional model training or predictor-based candidate selection was used, and the Unguided control differed only by selecting editing sites at random.

**Figure 6:**
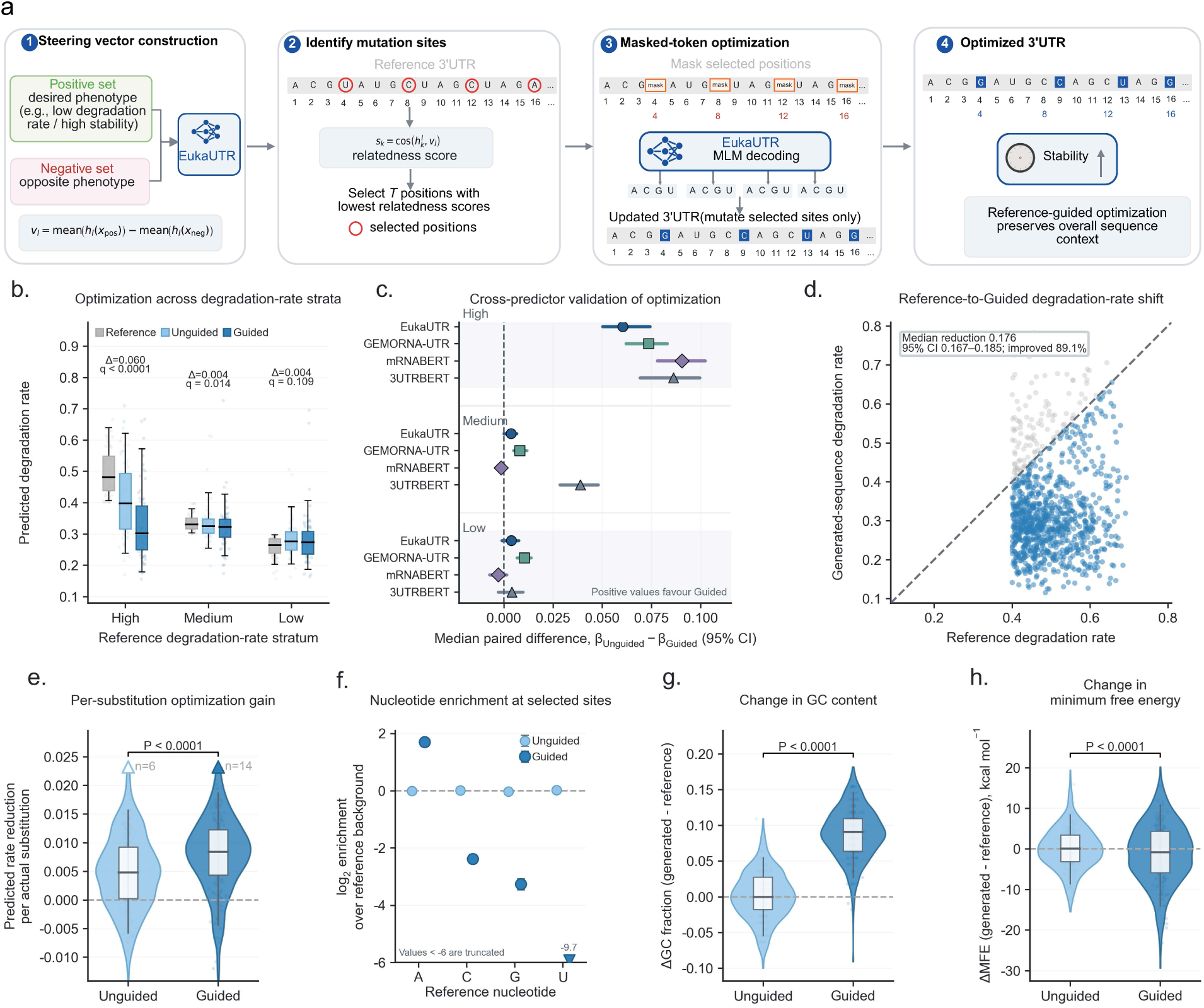
EukaUTR-Guide enables property-directed 3′ UTR editing for improved predicted mRNA stability. **a,** Overview of property-guided editing with EukaUTR-S2. **b,** Predicted degradation rates (*β*) for Reference, Unguided and Guided sequences stratified by the initial predicted degradation rate of the reference sequence. **c,** Cross-predictor comparison of predicted degradation rates between Guided and Unguided editing. **d,** Reference-to-Guided changes in predicted degradation rate for high-rate reference sequences. **e,** Predicted degradation-rate reduction per realized nucleotide substitution for Unguided and Guided editing. **f,** Nucleotide enrichment at positions selected by Unguided and Guided editing relative to reference-sequence composition. **g,h,** Changes in GC fraction (g) and predicted minimum free energy (MFE; **h**) after Unguided and Guided editing relative to the corresponding reference sequences. Error bars in **c,f** denote 95% bootstrap confidence intervals.

Optimization gains were greatest for sequences with high initial predicted degradation rates, which had the largest scope for improvement. Among high-rate references, the median predicted degradation rate decreased from 0.481 in Reference sequences to 0.397 after Unguided editing, a 17.5% reduction. Because Unguided editing selected sites randomly but used the same EukaUTR-S2 masked decoder, this reduction provides a baseline for context-aware decoding without property-guided site selection. Guided editing reduced the median rate further to 0.302, corresponding to a 37.2% reduction from Reference and a 23.9% reduction relative to Unguided editing (Holm-adjusted *P* < 0.0001; Fig. 6b). The additional benefit of Guided editing became smaller as the initial predicted degradation rate decreased, consistent with less room for further improvement in lower-rate sequences. The same pattern was supported by three external sequence models, all of which predicted lower degradation rates for Guided than for Unguided sequences in the high-rate group (Holm-adjusted *P* < 0.0001 for all comparisons; Fig. 6c and Supplementary Fig. 5a–c). The difference was smaller and less consistent in the medium-rate group and minimal in the low-rate group. Guided editing reduced the predicted degradation rate in 89.1% of high-rate reference sequences (Fig. 6d) and also produced a larger predicted reduction per realized nucleotide substitution than Unguided editing (*P* < 0.0001; Fig. 6e). Thus, the Guided effect was observed across most high-rate sequences and could not be explained simply by a greater number of nucleotide substitutions.

Guided editing preferentially selected nucleotide positions with a distinct base composition. Compared with the corresponding reference sequences, Guided-selected positions were enriched for A and depleted for C, G and especially U, whereas Unguided-selected positions closely matched the reference composition (Fig. 6f). Guided-selected sites were also more likely to change during decoding: only 8.5% retained the original nucleotide, compared with 35.2% of Unguided-selected sites (Supplementary Fig. 6a,b). At the sequence level, Guided editing increased GC content by about 9 percentage points, shifted the minimum free energy towards more negative values, and reduced predicted human miRNA 8-mer target-site abundance by approximately 58% relative to Reference sequences (Fig. 6g,h; Supplementary Fig. 7c).

Guided editing increased GC content by about 9 percentage points, a sequence feature associated with RNA stability [3, 34]. We therefore examined whether GC enrichment contributed to the predicted reduction in degradation rate. Changes in GC fraction were weakly correlated with predicted degradation-rate reduction for both Unguided and Guided editing (Supplementary Fig. 8b). Adjusting for ΔGC reduced, but did not eliminate, the estimated Guided effect in regression analyses, whereas no significant residual advantage was detected after matching sequences with similar ΔGC and initial predicted degradation rates (Supplementary Fig. 8a,c,d). Together, these analyses suggest that GC enrichment is an important component of the sequence changes underlying the predicted improvement produced by EukaUTR-Guide.

Together, these results show that EukaUTR can use a property-associated signal derived from a small number of sequences to guide targeted 3′ UTR editing without additional model training. EukaUTR-Guide shifted sequences towards lower predicted degradation rates, with consistent support across multiple sequence models. GC enrichment was an important sequence change associated with this effect, although experimental validation will be required to determine whether the predicted improvements increase mRNA stability in cells. EukaUTR-Guide therefore provides a framework for precise, property-guided 3′ UTR design.

## 3 Discussion

EukaUTR provides a unified framework for functional prediction, regulatory sequence analysis and design of eukaryotic 3′ UTRs. Its two-stage pre-training strategy combines broad eukaryotic sequence diversity with subsequent refinement on a high-confidence vertebrate corpus. Across 13 downstream tasks spanning post-transcriptional regulation, RNA stability and gene-expression outputs, all three EukaUTR models showed strong transfer performance. EukaUTR-S2 most often achieved the highest numerical performance, particularly for mRNA-stability and expression-output prediction, whereas EukaUTR-S2-LoRA performed best on several benchmarks, including eCLIP and APA. However, these gains were task-dependent, and direct comparisons among the three EukaUTR models did not consistently show significant differences. Together, these results suggest that Stage 2 adaptation can improve downstream transfer, but the extent of the benefit depends on both the task and the adaptation strategy.

The broad downstream performance suggests that EukaUTR captures sequence features relevant to multiple aspects of 3′ UTR regulation rather than features specific to a single prediction task. The local regulatory analyses provide further support for this interpretation. EukaUTR-S2 identified greater mutational sensitivity within the experimentally defined *CXCL2* ARE1 region and captured an AUA-rich sequence signal associated with RBMS3-bound regions. At the same time, prediction of individual nucleotide effects within ARE1 remained limited, and the small RBMS3 reporter panel did not show significant quantitative agreement between predicted and experimental effects. EukaUTR can therefore identify regional and sequence-context signals, but these results do not support nucleotide-level or mechanistic interpretation.

EukaUTR also extends 3′ UTR modelling from prediction to sequence generation and targeted design. De novo-generated sequences showed multiple natural-like properties, including nucleotide composition, local *k*-mer patterns, predicted structural properties and cis-regulatory-element frequencies, while showing no matches to the pre-training corpus at ≥ 80% sequence identity and ≥ 0.8 alignment coverage. These results support the sequence-level similarity and novelty of EukaUTR-generated sequences, although their biological activity remains to be established experimentally. EukaUTR-Guide further uses a property-associated signal derived from a small number of sequences to prioritize editing sites without additional model training or predictor-based selection of candidate substitutions. In the mRNA-stability example studied here, Guided editing shifted sequences towards lower predicted degradation rates, with the same direction of improvement supported by multiple sequence models.

Guided editing also produced systematic changes in sequence composition, including an increase in GC content, more negative RNAfold-predicted minimum free energy and fewer predicted human miRNA 8-mer target sites. GC enrichment is particularly notable because GC content has been associated with RNA stability [3, 34]. Sensitivity analyses indicated that this increase in GC content contributes importantly to the predicted improvement produced by EukaUTR-Guide. Rather than representing an independent validation of stability, these findings suggest that GC composition is one of the stability-associated sequence properties captured by the current editing strategy. Whether additional sequence features beyond GC content contribute to the predicted stability gains of EukaUTR-Guide remains to be determined.

Several limitations remain. Although EukaUTR was pretrained across broad eukaryotic sequence diversity, functional evaluation was concentrated primarily on human and zebrafish datasets. Testing across additional eukaryotic lineages will therefore be important for determining how broadly the learned sequence information transfers across species. In addition, both de novo-generated and EukaUTR-Guide-edited sequences have so far been evaluated computationally. Experimental measurements of mRNA stability, translation and other regulatory outputs will therefore be needed to determine whether the predicted properties of these sequences are retained in cells. More broadly, integrating broader experimental validation with property-guided sequence design may enable EukaUTR to support more precise design of 3′ UTRs for research and therapeutic applications.

## 4 Methods

### 4.1 Pre-training

#### Pre-training datasets

The Stage 1 and Stage 2 pre-training corpora were constructed from Ensembl and Ensembl Genomes annotations [20, 21]. Stage 1 comprised annotated 3′ UTRs spanning major eukaryotic lineages, including vertebrates, non-vertebrate metazoans, plants, fungi and protists. Stage 2 comprised a curated high-confidence corpus from human, mouse, rat and zebrafish, all four of which were excluded from Stage 1. Sequences shorter than 32 nt or longer than 10,000 nt were removed, and redundancy was reduced by MMseqs2 clustering at 80% sequence identity [22]. The resulting Stage 1 corpus contained 7,025,323 non-redundant 3′ UTRs from 1,715 genomes (3.268 billion nucleotides), whereas Stage 2 contained 59,221 non-redundant 3′ UTRs (48.20 million nucleotides).

#### Model architecture and sequence processing

EukaUTR uses an ESM-2-style Transformer encoder [23–25] comprising 12 layers, 12 self-attention heads per layer and a hidden dimension of 768, with approximately 87 million parameters. Sequences were converted to the RNA alphabet and tokenized at single-nucleotide resolution, with up to 1,022 nt represented in each input. During pre-training, longer sequences were split into overlapping windows with a 200-nt overlap rather than truncated, ensuring that all nucleotide positions were retained for pre-training. Additional details on the vocabulary, model architecture and windowing procedure are provided in the Supplementary Note.

#### Pre-training and model refinement

EukaUTR was pretrained using a masked language-modelling objective. In Stage 1, 15% of nucleotide positions were selected for prediction; of these, 80% were replaced with the mask token, 10% with a random nucleotide and 10% were left unchanged. We reserved 1% of the broad eukaryotic corpus for validation and optimized the model using AdamW [35] with a peak learning rate of 4 × 10^−4^. The checkpoint with the lowest validation loss was retained as EukaUTR-S1.

Starting from EukaUTR-S1, we further refined the model on the high-confidence Stage 2 corpus using two adaptation strategies with the same masked language-modelling objective. Full-parameter continued pre-training updated all model parameters with a learning rate of 1 × 10^−4^, yielding EukaUTR-S2. In parallel, LoRA-based adaptation [26] (*r* = 8, *α* = 16) introduced approximately 1.3 million trainable parameters, corresponding to about 1.5% of the full model, and yielded EukaUTR-S2-LoRA. We reserved 5% of the Stage 2 corpus for validation. Additional details on the optimization schedules, LoRA insertion sites, batch construction, mixed-precision training, hardware and checkpoint selection are provided in the Supplementary Note.

### 4.2 Downstream task evaluation

#### Downstream datasets and prediction tasks

We evaluated EukaUTR across 13 prediction tasks derived from nine published datasets. These included established benchmark datasets as well as experimental datasets that we processed to define prediction targets, spanning RBP binding, m^6^A modification, alternative polyadenylation, RNA degradation and half-life, RNA abundance, translational efficiency, protein output and mean ribosome load (Supplementary Table 3).

RBP-binding prediction was evaluated in two settings: a fivefold cross-validated eCLIP benchmark spanning 22 RBP–cell-line combinations and a gene-held-out CLIP benchmark comprising 31 experiments, in which test sequences were derived from genes excluded from downstream model fitting [17]. m^6^A-site prediction was evaluated across nine human cell lines using the benchmark compiled from m^6^A-Atlas [17, 36]. Alternative polyadenylation was evaluated as quantitative prediction of proximal polyadenylation-site usage using the BEACON benchmark, which is based on the APARENT massively parallel reporter assay [29, 30].

mRNA degradation and half-life were evaluated using three datasets spanning reporter-based and cellular measurements. The zebrafish UTR-seq dataset of Rabani et al. [2] was used to define two prediction tasks: experimentally inferred degradation rate (*β*) and the corresponding half-life, calculated as *t*_1_*_/_*_2_ = ln(2)*/β* for sequences with valid positive *β* estimates. The fast-UTR dataset of Zhao et al. [9] provided experimentally estimated half-life for human reporter mRNAs. For cellular RNA decay, we constructed a benchmark from the SLAM-seq dataset of Karner et al. [32]. Because the released processed data did not contain the final kinetic decay estimates, we reconstructed a relative decay phenotype from nucleotide-conversion signal and steady-state RNA abundance and averaged this phenotype across six breast cancer cell lines.

Gene-expression outputs were evaluated using the full-length human 3′ UTR reporter dataset of West et al. [3] and the viromics MPRA dataset of Seo et al. [33]. In the West et al. dataset, barcoded 3′ UTRs were assayed downstream of a GFP reporter under CAG and PGK promoter contexts, with protein output, RNA abundance and translational efficiency used as separate prediction targets. Performance was evaluated separately for the two promoter contexts, and the corresponding scores were averaged for presentation in the main text. In the Seo et al. dataset, 130-nt viral genomic tiles were inserted into the 3′ UTR of a common reporter and evaluated for RNA abundance and mean ribosome load. Cross-validation was grouped by viral genome accession so that all tiles from the same viral genome were assigned to the same fold, preventing overlapping tiles from appearing in both training and held-out partitions.

Dataset-specific sample sizes and sequence lengths are summarized in Supplementary Table 3, with sequence processing, inclusion and exclusion criteria, label construction and normalization procedures described in the Supplementary Note.

#### Fine-tuning and evaluation

EukaUTR-S1, EukaUTR-S2 and EukaUTR-S2-LoRA were evaluated as three model configurations representing Stage 1 pre-training, full-parameter Stage 2 refinement and LoRA-based Stage 2 refinement, respectively. All three configurations were carried forward to downstream evaluation to assess the effects of Stage 2 refinement and adaptation strategy.

Original benchmark partitions were retained whenever available. The eCLIP benchmark used fivefold cross-validation, the CLIP benchmark retained its gene-held-out split, the m^6^A benchmark followed the original ten balanced evaluations for each cellular context, and BEACON [30] retained its predefined splits. The remaining regression tasks used tenfold cross-validation. For the viral MPRA benchmark, folds were grouped by viral genome accession so that all tiles from the same viral genome remained within the same fold. Identical task-specific partitions were used across all retrained models.

For each task, retrained language models were coupled to a lightweight prediction head using the final-layer <cls> representation and fine-tuned end-to-end, unless otherwise specified. To ensure comparable inputs across models, sequences were restricted to a maximum of 1,022 nt; longer sequences were truncated to the 5′-proximal 1,022 nt for all models. This length restriction was applied only during downstream benchmarking. Checkpoints were selected using validation performance before evaluation on held-out data. Classification tasks were evaluated primarily using F1 score and Matthews correlation coefficient (MCC), whereas regression tasks were evaluated using Pearson and Spearman correlations. Detailed training hyperparameters and additional evaluation metrics are provided in the Supplementary Note.

### 4.3 Local cis-regulatory analyses

#### CXCL2 saturation-mutagenesis analysis

Local mutational effects were analysed using saturation-mutagenesis data from a 67-nt region of the human *CXCL2* 3′ UTR comprising the 21-nt ARE1 element and 46 nt of flanking sequence [9]. EukaUTR-S2 was fine-tuned on the separate fast-UTR mRNA half-life benchmark and used to score the wild-type sequence and all 201 possible single-nucleotide substitutions. The *CXCL2* wild-type sequence and all corresponding mutants were excluded from model training, validation and checkpoint selection. Predicted mutational effects were defined as the mutant-minus-wild-type change in predicted mRNA half-life and were compared with experimentally measured changes in steady-state reporter mRNA abundance relative to wild type. Regional mutational sensitivity was assessed by comparing the absolute predicted effect magnitudes between substitutions within ARE1 and those in the flanking sequence. Detailed sequence processing, prediction aggregation and statistical procedures are provided in the Supplementary Note.

#### RBMS3-associated sequence-context analysis

RBMS3-associated sequence signals were analysed using reporter and CLIP-seq data from Karner et al. [32]. EukaUTR-S2 was fine-tuned on the Rabani et al. zebrafish UTR-seq benchmark [2] to predict mRNA degradation rate and was used for all sequence scoring. In the reporter analysis, 13 RBMS3-bound reference elements and their dinucleotide-preserving scrambled controls were scored separately, and predicted paired effects were defined as the scrambled-minus-reference difference in predicted degradation rate. These effects were compared with the corresponding experimentally measured reference-versus-scrambled reporter effects.

For the endogenous sequence-context analysis, 101-nt windows centred on RBMS3 CLIP sites were paired with non-overlapping control windows from the same 3′ UTR, yielding 11,248 matched pairs. AUA abundance and predicted degradation scores were compared between RBMS3-bound and matched-control windows. An interaction model was further used to test whether the association between AUA density and predicted degradation differed between the two window classes. Detailed sequence construction, control matching and statistical procedures are provided in the Supplementary Note.

### 4.4 De novo 3**′** UTR generation and benchmarking

#### Generation benchmark and comparator models

Natural 3′ UTRs between 100 and 1,022 nt were used to define the reference length distribution. A target-length vector of 10,000 sequences was sampled once from this distribution and reused across all generation methods, providing one-to-one length matching. An independent natural set matched to the same target-length vector was used as the reference.

EukaUTR-S1, EukaUTR-S2 and EukaUTR-S2-LoRA generated sequences from fully masked inputs using iterative confidence-guided decoding for up to 20 rounds. At each round, remaining masked positions were ranked by model confidence, a linearly scheduled subset of the highest-confidence positions was selected, and nucleotides were sampled from the model probabilities over A, C, G and U at temperature 1.0. EukaUTR was compared with GEMORNA-UTR [18], 3UTRBERT [17] and mRNABERT [19]. Because 3UTRBERT and mRNABERT do not provide native de novo generation procedures, they were evaluated using the same iterative masked-decoding framework with model-specific tokenization and vocabulary handling, whereas GEMORNA-UTR was generated using its native generative procedure. No natural reference sequence or downstream functional predictor was used during EukaUTR generation. Detailed decoding procedures are provided in the Supplementary Note.

#### Sequence-level and embedding-based evaluation

Generated sequences were compared with the length-matched natural reference using measures of nucleotide composition, predicted RNA structure, local sequence composition and regulatory-element content. These included GC content, length-normalized RNAfold minimum free energy, *k*-mer frequency correlations for *k* = 3–6, densities of canonical polyadenylation signals and other post-transcriptional regulatory motifs, predicted human miRNA 8-mer target sites and homopolymer statistics. Sequence naturalness was additionally quantified using EukaUTR-S2 as a fixed masked-language-model evaluator, based on the exponentiated mean conditional log-probability of each nucleotide when masked individually. Because EukaUTR-S2 was also evaluated as a generator, this score was treated as a model-based measure rather than an independent measure of biological function.

Embedding-level similarity to the natural reference was assessed using EukaUTR-S2 <cls> embeddings and quantified by Fréchet distance and squared maximum mean discrepancy (MMD^2^) [37, 38]. As these metrics were computed in EukaUTR-S2 embedding space, they were interpreted as internal measures of distributional similarity. Metric definitions, motif patterns, RNAfold settings, embedding extraction and MMD parameters are provided in the Supplementary Note.

#### Nearest-neighbour sequence similarity

Sequence similarity was evaluated using MMseqs2 [22] both within each generated set and against the complete EukaUTR pre-training corpus. Self-matches were excluded from within-set searches. Sequences with a nearest-neighbour match at ≥ 80% sequence identity and ≥ 0.8 alignment coverage were classified as having a high-similarity near-neighbour, and frequencies were reported per 10,000 evaluated sequences. Detailed MMseqs2 search parameters are provided in the Supplementary Note.

### 4.5 EukaUTR-Guide implementation and evaluation

EukaUTR-Guide uses a property-associated direction derived from EukaUTR representations to prioritize editing sites, followed by masked-nucleotide decoding with EukaUTR-S2. Inspired by steering-vector approaches to protein sequence design [39], we defined a low-degradation steering direction from EukaUTR-S2 representations of sequences with contrasting predicted degradation rates. This direction was used to prioritize editing positions.

Steering and reference sets were constructed from 110-nt fragments sampled from 3′ UTRs of highly expressed human genes. Fragments were scored using an EukaUTR-S2 degradation-rate predictor fine-tuned on the Rabani et al. UTR-seq benchmark [2]. One hundred fragments with low predicted degradation rates and 100 with high predicted degradation rates were selected to define the desired and contrasting representation states, respectively. Steering fragments were excluded from the reference sets, which were stratified by their initial predicted degradation rates into high (*β̂* > 0.4), medium (0.3 ≤ *β̂* ≤ 0.4) and low (*β̂* < 0.3) groups.

For Guided editing, nucleotide positions were ranked by their cosine similarity to the low-degradation representation direction. In each round, the four least-aligned positions were masked and decoded by EukaUTR-S2 using multinomial sampling over A, C, G and U. Editing was repeated for eight rounds, with sequence representations and position rankings recalculated after each round. An Unguided control selected four nucleotide positions at random in each round and used the same EukaUTR-S2 decoder, sampling procedure and number of editing rounds. Guided and Unguided editing therefore differed only in how candidate editing positions were selected.

Reference, Guided and Unguided sequences were evaluated using the EukaUTR-S2 degradation-rate predictor and degradation-rate predictors based on GEMORNA-UTR, mRNABERT and 3UTRBERT, each fine-tuned on the same Rabani et al. benchmark. Predicted degradation rates were used to construct the steering sets and to evaluate final editing outcomes, but were not used during decoding to accept, reject or rank candidate nucleotide substitutions. Detailed procedures for steering-set construction, representation-direction estimation, decoding, sequence-property analysis and statistical evaluation are provided in the Supplementary Note.

## Code availability

The latest version of the code is available at https://github.com/meilanglang/EukaUTR under the MIT license. The pre-trained model checkpoints are available at https://huggingface.co/langmei/EukaUTR/.

### Data availability

The processed Stage 1 and Stage 2 pre-training corpora and the downstream benchmark datasets used in this study are available through Zenodo at https://doi.org/10.5281/zenodo.21790090. The deposited materials include the pre-training sequences, processed downstream-task datasets and associated metadata required to reproduce the analyses. The original source data remain available from the databases and studies cited in the Methods and Supplementary Note.

## Acknowledgements

This work was supported by the Zhejiang Provincial Science and Technology Program (2025C01129 to X.L.) and the China Postdoctoral Science Foundation (2025M772773 to Z.W.).

## Author contributions

X.L., K.Y.T., and J.Z. supervised the study and provided resources. M.L. and X.L. conceptualized the study, designed the EukaUTR framework and model architecture, implemented the computational methods, performed model training and data analyses, and wrote the original draft of the manuscript. X.F. contributed to downstream-task data processing and validation and to the computational evaluation of de novo-generated sequences. M.C. performed LoRA-based model fine-tuning. All authors reviewed and edited the manuscript.

## Supplementary Information

### Supplementary Note

#### Construction of the Stage 1 and Stage 2 pre-training corpora

##### Stage 1 broad eukaryotic corpus

The Stage 1 corpus was constructed from Ensembl and Ensembl Genomes annotations [20, 21] to capture broad 3′ UTR sequence diversity across the major eukaryotic divisions represented in the Ensembl ecosystem. Vertebrate annotations were obtained from Ensembl release 114, whereas annotations for non-vertebrate Metazoa, Plants, Fungi and Protists were obtained from Ensembl Genomes release 61.

For each genome, the corresponding genome assembly and gene-annotation files were downloaded and parsed to extract transcript regions explicitly annotated as 3′ UTRs. Genomes were excluded when 3′ UTR annotations could not be parsed, transcript structures were incomplete, genomic coordinates were invalid or no valid 3′ UTR sequence could be recovered after coordinate validation. Ensembl Bacteria was examined but excluded because bacterial and archaeal annotations generally do not define canonical eukaryotic 3′ UTR features suitable for this analysis.

Before sequence filtering and redundancy reduction, the Stage 1 candidate pool contained 26,242,237 3′ UTR sequences from 1,748 genomes, comprising approximately 18.787 billion nucleotides. The source genomes included 325 vertebrates, 354 non-vertebrate metazoans, 199 plants, 740 fungi and 130 protists. Vertebrates contributed 8,275,287 candidate sequences, Plants contributed 7,672,967, Metazoa contributed 6,715,739, Fungi contributed 3,223,145 and Protists contributed 355,099.

Sequences shorter than 32 nt or longer than 10,000 nt were removed. The remaining sequences were clustered using MMseqs2 [22] at an 80% sequence-identity threshold, and one representative sequence was retained from each cluster.

The final Stage 1 corpus contained 7,025,323 non-redundant 3′ UTRs from 1,715 genomes, comprising approximately 3.268 billion nucleotides. It included 319 vertebrate genomes, 353 metazoan genomes, 198 plant genomes, 719 fungal genomes and 126 protist genomes. The overall mean and median sequence lengths were 465 and 234 nt, respectively.

Metazoa contributed 2,474,116 sequences comprising 1.386 billion nucleotides, Fungi contributed 1,973,341 sequences comprising 0.476 billion nucleotides, Plants contributed 1,308,367 sequences comprising 0.528 billion nucleotides, Vertebrates contributed 1,089,644 sequences comprising 0.821 billion nucleotides and Protists contributed 179,855 sequences comprising 0.057 billion nucleotides. Mean sequence lengths were 753.21 nt for Vertebrates, 560.40 nt for Metazoa, 403.77 nt for Plants, 241.24 nt for Fungi and 315.79 nt for Protists. The corresponding median lengths were 464, 248, 267, 168 and 153 nt.

##### Stage 2 high-confidence corpus

The Stage 2 corpus was constructed independently from curated Ensembl transcript annotations for human (*Homo sapiens*), mouse (*Mus musculus*), rat (*Rattus norvegicus*) and zebrafish (*Danio rerio*), four species for which Ensembl integrates automated gene annotation with HAVANA manual annotation. These four species were excluded from Stage 1 to preserve species-level separation between the two pre-training stages.

Before redundancy reduction, the Stage 2 candidate pool contained 213,579 3′ UTR sequences comprising approximately 0.252 billion nucleotides. The same length filters and MMseqs2-based redundancy-reduction procedure used for Stage 1 were applied.

The final Stage 2 corpus contained 59,221 non-redundant 3′ UTRs comprising 48.20 million nucleotides. The mean sequence length was 813.88 nt and the median was 421 nt. Human contributed 20,677 sequences comprising 18.81 million nucleotides, with mean and median lengths of 909.68 and 418 nt. Mouse contributed 14,355 sequences comprising 13.02 million nucleotides, with mean and median lengths of 907.03 and 468 nt. Rat contributed 9,867 sequences comprising 7.80 million nucleotides, with mean and median lengths of 790.55 and 466 nt. Zebrafish contributed 14,322 sequences comprising 8.57 million nucleotides, with mean and median lengths of 598.28 and 373 nt.

Together, the two stages comprised 7,084,544 non-redundant sequences from 1,719 unique genomes. Stage 1 provided broad phylogenetic coverage, whereas Stage 2 provided a smaller curated vertebrate corpus for continued pre-training.

#### EukaUTR architecture and pre-training

##### Model architecture and vocabulary

EukaUTR was implemented using an ESM-2-style Transformer encoder. The model comprised 12 Transformer layers, 12 self-attention heads per layer and a hidden dimension of 768, with approximately 87 million parameters. Dropout, attention dropout and activation dropout were each set to 0.1. The maximum input length was 1,024 tokens, corresponding to up to 1,022 nucleotide tokens together with the <cls> and <eos> tokens.

Sequences were converted to uppercase and standardized to the RNA alphabet by replacing T with U. Ambiguous or unsupported nucleotide symbols, including R, Y, K, M, S, W, B, D, H, V and N, were represented as X. Sequences were tokenized at single-nucleotide resolution using the vocabulary

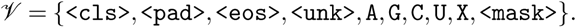

##### Processing of long sequences

Sequences of up to 1,022 nt were processed as single inputs. Longer sequences were divided into overlapping windows of up to 1,022 nt, with adjacent windows sharing 200 nt. Each window was treated as an independent pre-training instance. If the terminal segment was shorter than the maximum window length, it was retained as the final window rather than discarded. This procedure allowed all nucleotide positions in long 3′ UTRs to be included in pre-training while preserving overlapping sequence context between adjacent windows.

##### Masked language-modelling objective

Both pre-training stages used the same masked language-modelling objective. For each training instance, 15% of nucleotide positions were selected as prediction targets. Of these positions, 80% were replaced with <mask>, 10% with a nucleotide sampled from {A, C, G, U}, and 10% were left unchanged. The training loss was the mean cross-entropy over the selected prediction positions.

##### Stage 1 pre-training

Stage 1 pre-training was performed on the broad eukaryotic corpus, with 1% of the training data reserved for validation. Model parameters were optimized using AdamW with a peak learning rate of 4 × 10^−4^. The learning rate was increased linearly during the first 10% of optimization steps and then decayed linearly to 10% of the peak learning rate.

Training was performed using distributed data parallelism on four NVIDIA A100 GPUs. The checkpoint with the lowest validation loss was retained as EukaUTR-S1 and used to initialize Stage 2.

##### Stage 2 full-parameter continued pre-training

For full-parameter Stage 2 continued pre-training, EukaUTR-S1 was further trained on the high-confidence Stage 2 corpus, with all model parameters updated. Five per cent of the Stage 2 data were reserved for validation. The learning rate was warmed up linearly to 1 × 10^−4^ over the first 500 optimization steps and then decayed linearly to zero.

Training was performed using distributed data parallelism on two NVIDIA A100 GPUs. The checkpoint with the lowest validation loss was retained as EukaUTR-S2.

##### Stage 2 LoRA continued pre-training

For parameter-efficient Stage 2 continued pre-training, the EukaUTR-S1 backbone was frozen and low-rank adaptation (LoRA) modules were inserted into six linear projections in each Transformer layer: the query, key, value and output projections of the self-attention module and the two feed-forward projections.

The final LoRA configuration used rank *r* = 8, *α* = 16 (*α/r* = 2), a LoRA dropout of 0.05 and no trainable bias terms. This configuration introduced approximately 1.3 million trainable parameters, corresponding to approximately 1.5% of the parameter count of the full model.

LoRA parameters were optimized using AdamW. The learning rate was warmed up linearly to 3 × 10^−5^ over the first 500 optimization steps and then decayed linearly to zero. Early stopping was applied with a patience of 10 epochs and a minimum validation-loss improvement of 10^−4^. Training was performed on two NVIDIA A100 GPUs, and the checkpoint with the lowest validation loss was retained as EukaUTR-S2-LoRA.

##### Batch construction and numerical precision

Stage 1 and both Stage 2 training procedures used FP16 mixed-precision training and dynamic length-based bucketing. Sequences of similar lengths were grouped into batches containing at most 32,768 nucleotide tokens, reducing padding while maintaining a fixed upper bound on the token-level batch size. During distributed training, batches were shuffled at each epoch and partitioned across training processes.

#### Downstream datasets, fine-tuning and evaluation

We evaluated EukaUTR across 13 prediction tasks derived from nine source datasets, including established benchmarks and experimental datasets that were processed to define prediction targets. The tasks covered RBP binding, m^6^A modification, alternative polyadenylation, mRNA degradation and half-life, RNA abundance, translational efficiency, protein output and mean ribosome load. Original benchmark partitions were retained whenever available; for the remaining tasks, fixed cross-validation partitions were constructed once and reused across all retrained models.

##### eCLIP RBP-binding prediction

The eCLIP RBP-binding benchmark was obtained from the processed dataset released with 3UTRBERT [17] and was derived from human ENCODE eCLIP experiments in K562 and HepG2 cells [40]. The benchmark comprised 552,682 100-nt sequences across 22 RBP–cell-line combinations. Sequence redundancy had been reduced using CD-HIT-EST at 80% sequence identity, and each task contained approximately one positive sequence for every two negative sequences. The original fivefold cross-validation protocol was retained, with identical folds used across all retrained models.

##### Gene-held-out CLIP RBP-binding prediction

The gene-held-out CLIP benchmark was also obtained from the processed 3UTRBERT dataset [17]. It comprised 120,881 101-nt sequences from 31 CLIP experiments spanning 19 RBPs. Positive sequences were centred on high-confidence CLIP sites, whereas negative sequences were sampled from genes without detected interactions. The predefined gene-held-out split was retained so that genes represented in the held-out partition were absent from downstream model fitting.

##### Human m^6^A-site prediction

The human m^6^A benchmark was obtained from the processed 3UTRBERT dataset [17], with experimentally supported sites derived from m^6^A-Atlas [36]. The dataset comprised 928,026 41-nt sequences centred on candidate adenosines across nine human cell lines. High-confidence m^6^A sites were used as positive examples, and non-annotated adenosines from the same transcripts were used as candidate negatives. Following the original protocol, the negative examples were divided into ten subsets, each of which was paired with the corresponding positive set to produce ten approximately balanced evaluations per cellular context.

##### Alternative polyadenylation prediction

The alternative polyadenylation benchmark was obtained from BEACON [30] and was derived from the APARENT massively parallel reporter assay [29]. It comprised 228,388 186-nt sequences, with proximal polyadenylation-site usage as the quantitative prediction target. The predefined training, validation and test partitions were retained without modification.

##### Zebrafish reporter degradation-rate and half-life prediction

Two prediction tasks were defined from the zebrafish UTR-seq reporter dataset of Rabani et al. [2]. The degradation-rate task comprised 90,000 110-nt reporter sequences with experimentally inferred degradation rates (*β*). For the 89,143 sequences with valid positive *β* estimates, the corresponding half-life was calculated as *t*_1_*_/_*_2_ = ln(2)*/β* and used as a second prediction target.

Both tasks used fixed tenfold cross-validation partitions shared across models. Within each fold, validation data used for checkpoint selection were drawn only from the corresponding training portion.

##### Human fast-UTR half-life prediction

The human reporter half-life benchmark was derived from the fast-UTR assay of Zhao et al. [9]. The processed dataset contained 1,967 160-nt reporter sequences with mRNA half-lives estimated from doxycycline-mediated transcriptional shutoff experiments. Performance was evaluated using fixed tenfold cross-validation partitions, with validation data drawn only from the training portion of each fold.

##### Breast cancer SLAM-seq mRNA decay prediction

A cellular RNA-decay benchmark was constructed from the SLAM-seq dataset of Karner et al. [32] (GEO GSE301535), comprising HCC1806, HCC38, MCF7, MDA-MB-231, MDA-MB-453 and ZR-75-1 cells. Gene-level nucleotide-conversion ratios were used as the metabolic-labelling signal, and steady-state RNA abundance was quantified from the corresponding RNA-sequencing data using Salmon and GENCODE v28.

Because the released processed data did not contain the final kinetic decay estimates used in the original study, we reconstructed a relative decay phenotype as the log_2_ ratio of nucleotide-conversion signal to steady-state RNA abundance, using pseudocounts of 10^−6^ for the conversion ratio and 1 for TPM. Conversion ratios ≥ 0.5 were treated as missing values, and genes with mean expression below 1 TPM were excluded. Replicates were first averaged within each cell line and then across the six cell lines to obtain a single cross-context target for each gene.

Each gene-level target was linked to a representative protein-coding transcript with an annotated 3′ UTR in GENCODE v28. After sequence filtering and redundancy reduction at 80% sequence identity, 4,918 non-redundant 3′ UTRs were retained. During model evaluation, sequences longer than 1,022 nt were represented by their 5′-proximal 1,022 nt. Fixed tenfold cross-validation partitions were used across all retrained models. The resulting target should therefore be interpreted as a reconstructed relative mRNA-decay phenotype rather than an absolute kinetic degradation rate.

##### Full-length human 3′ UTR reporter prediction

The full-length human 3′ UTR reporter benchmark was derived from West et al. [3]. Barcoded 3′ UTRs were assayed downstream of a GFP reporter under CAG and PGK promoter contexts. Protein output, RNA abundance and translational efficiency were treated as separate quantitative prediction targets.

After redundancy reduction at 80% sequence identity, 1,343 non-redundant 3′ UTRs were retained. During model evaluation, sequences longer than 1,022 nt were represented by their 5′-proximal 1,022 nt, whereas the prediction targets remained those measured from the original full-length reporter constructs. Each target was modelled separately under the CAG and PGK promoter contexts using identical tenfold cross-validation partitions across models. Main-text performance values were calculated as the arithmetic mean of the corresponding CAG-and PGK-specific scores.

##### Viromics MPRA prediction

The viromics MPRA benchmark was derived from Seo et al. [33]. It comprised 30,155 130-nt viral genomic tiles assayed after insertion into the 3′ UTR of a common reporter. RNA abundance was quantified from RNA reads normalized to the corresponding DNA reads, and mean ribosome load (MRL) was obtained from polysome-profiling measurements. The two measurements were treated as separate quantitative prediction targets.

Because the library contained overlapping genomic tiles, tenfold cross-validation was grouped by viral genome accession so that all tiles from the same accession remained within the same fold. This prevented overlapping tiles from the same viral genome from appearing in both training and held-out partitions.

##### Downstream fine-tuning and evaluation

All retrained language models were coupled to a lightweight prediction head using the final-layer <cls> representation and were fine-tuned end-to-end. The prediction head used a dropout rate of 0.25. AdamW optimization used learning rates of 1 × 10^−4^ for the encoder and 4 × 10^−4^ for the prediction head, with 10% linear warm-up followed by cosine decay and gradient clipping at a norm of 1.0.

To ensure comparable sequence inputs across models, all downstream inputs were restricted to a maximum of 1,022 nt. Sequences exceeding this limit were represented by their 5′-proximal 1,022 nt for all models. This restriction was applied only during downstream benchmarking.

Regression tasks used Huber loss, except for alternative polyadenylation prediction, which used mean-squared error; classification tasks used cross-entropy loss. Models were trained for up to 20–50 epochs with task-specific batch sizes and early stopping. Checkpoints were selected using validation performance before evaluation on the corresponding held-out data. Classification performance was assessed primarily using F1 score and Matthews correlation coefficient (MCC), whereas regression performance was assessed using Pearson and Spearman correlations.

##### Statistical comparison of downstream models

For tenfold cross-validation benchmarks, statistical comparisons were based on paired scores from the same ten held-out folds and were performed using two-sided Wilcoxon signed-rank tests. For comparisons between the three EukaUTR models and the external baselines, Holm correction was applied separately within each prediction task and evaluation metric across the nine EukaUTR-versus-baseline comparisons. Comparisons among EukaUTR-S1, EukaUTR-S2 and EukaUTR-S2-LoRA were corrected separately across the three pairwise comparisons within each prediction task and evaluation metric.

#### Local cis-regulatory analyses

##### *CXCL2* ARE1 saturation-mutagenesis analysis

The *CXCL2* saturation-mutagenesis dataset was obtained from the fast-UTR reporter assay of Zhao et al. [9]. The analysed region comprised 67 nt of the human *CXCL2* 3′ UTR (positions 138–204 in the reporter construct), including the experimentally characterized 21-nt ARE1 element and 46 nt of flanking sequence. Substitution of each nucleotide by the three alternative canonical nucleotides yielded 201 single-nucleotide mutants.

Experimental mutational effects were defined as changes in steady-state reporter mRNA abundance relative to the wild-type construct. Reporter abundance in the original assay was quantified from RNA read counts normalized to the corresponding genomic-DNA counts [9]. Positive experimental effects therefore indicate increased steady-state mRNA abundance relative to wild type.

Model predictions were obtained using an EukaUTR-S2 half-life predictor fine-tuned on the separate fast-UTR mRNA half-life benchmark from Zhao et al. The wild-type *CXCL2* sequence and all 201 corresponding mutants were excluded from the data used for model training, validation and checkpoint selection. The wild-type sequence and each mutant were scored using the ten fold-specific models obtained during cross-validation. For each model, the predicted mutational effect was defined as the mutant-minus-wild-type change in predicted mRNA half-life, and the final effect for each substitution was calculated as the mean across the ten models. Positive predicted effects therefore indicate a longer predicted half-life for the mutant than for the wild-type sequence.

Predicted and experimental effects were compared across all 201 substitutions using two-sided Pearson and Spearman correlation tests. Regional mutational sensitivity was assessed by comparing the absolute predicted effect magnitudes of substitutions within ARE1 with those in the flanking sequence using a two-sided Wilcoxon rank-sum test. Within ARE1, nucleotide-level agreement was assessed separately using Pearson and Spearman correlations and by comparing the upper 20% of substitutions ranked by absolute predicted and experimental effect magnitude. For visualization, experimental and predicted effects were arranged as position-by-mutant-nucleotide matrices, with the wild-type nucleotide at each position left blank.

##### RBMS3-bound 3′ UTR reporter analysis

The RBMS3 reporter dataset was obtained from Karner et al. [32] (GEO GSE301428). Thirteen RBMS3-bound 3′ UTR reference elements (REF) and corresponding dinucleotide-preserving scrambled controls (SCR) were assayed in MDA-MB-231 cells using a dual-reporter system.

RNA and DNA barcode counts were normalized by library size. Experimental REF–SCR activity was calculated as the log_2_ ratio of REF-to-SCR RNA-to-DNA signal and averaged across control (*shCTRL*) replicates. Positive values therefore indicate greater reporter activity for REF than for SCR.

REF and SCR sequences were scored using an EukaUTR-S2 degradation-rate predictor fine-tuned on the Rabani et al. zebrafish UTR-seq benchmark [2]. The predicted paired effect was calculated as the SCR-minus-REF difference in predicted degradation rate, such that positive values indicate lower predicted degradation for REF. Predicted and experimental effects were compared using Pearson and Spearman correlations, and directional concordance was defined as agreement in effect sign. Given the small number of pairs (*n* = 13), this analysis was used primarily to assess directional agreement.

##### RBMS3 CLIP-site sequence-context analysis

RBMS3 CLIP-seq data were obtained from Karner et al. [32] (GEO GSE301429). After mitochondrial sites were removed, 27,486 RBMS3 cross-link-induced mutation sites (CIMS) remained. Strand-specific intersection with protein-coding 3′ UTR annotations from GENCODE v28 identified 20,627 sites within annotated 3′ UTRs, corresponding to 6,157 genes.

A 101-nt window centred on each overlapping RBMS3 CIMS site was extracted. Windows that were not fully contained within an annotated 3′ UTR were excluded, leaving 11,613 eligible RBMS3-site windows. For each retained window, a non-overlapping 101-nt control window was sampled from the same annotated 3′ UTR while excluding regions overlapping RBMS3 CIMS sites. This procedure yielded 11,248 matched RBMS3-bound–control pairs.

AUA abundance was quantified in each window as both the number of AUA trinucleotides and their density. The same windows were scored using the EukaUTR-S2 degradation-rate predictor. AUA abundance and predicted degradation rates were compared between matched RBMS3-bound and control windows using two-sided paired Wilcoxon signed-rank tests.

To determine whether the association between AUA density and predicted degradation differed between RBMS3-bound and control windows, standardized predicted degradation rates were regressed on window class, standardized AUA density and their interaction. The interaction term tested whether the AUA–predicted-degradation association differed between the two window classes. Cluster-robust standard errors were calculated at the matched-pair level.

Because EukaUTR receives sequence alone and does not model RBMS3 abundance, binding occupancy or cellular state, these predicted degradation rates were interpreted as sequence-derived signals associated with RBMS3-bound regions, rather than as estimates of the causal effect of RBMS3 binding on mRNA degradation.

#### De novo 3**′** UTR generation and benchmarking

##### Benchmark construction and target-length matching

Natural 3′ UTRs from the complete EukaUTR pre-training corpus with lengths between 100 and 1,022 nt were used to define the empirical target-length distribution. A vector of 10,000 target lengths was sampled once from this distribution and reused across all generation methods and the random control, providing one-to-one length matching. An independent set of natural 3′ UTRs matched to the same target-length vector was used as the natural reference. Random-control sequences were generated by independently sampling A, C, G and U with equal probability at each nucleotide position.

##### De novo generation with EukaUTR

EukaUTR-S1, EukaUTR-S2 and EukaUTR-S2-LoRA were used for template-free de novo generation from fully masked inputs of the predefined target lengths. Generation proceeded for up to 20 iterative decoding rounds. At each round, probabilities at the remaining masked positions were restricted to A, C, G and U. Positions were ranked by the maximum predicted nucleotide probability, and a linearly scheduled subset of the highest-confidence positions was decoded by multinomial sampling at temperature 1.0 without top-*p* truncation. The updated sequence was then used for the next round until all masked positions were resolved. Generated sequences were required to contain only canonical RNA nucleotides and to match their predefined target lengths exactly. No natural reference sequence, downstream functional predictor or predictor-based acceptance–rejection procedure was used during EukaUTR generation.

##### Comparator generation

EukaUTR-generated sequences were compared with GEMORNA-UTR, 3UTRBERT and mRNABERT using publicly available model weights. Because 3UTRBERT and mRNABERT do not provide native de novo generation procedures, they were evaluated using the same iterative confidence-guided masked-decoding framework, with model-specific tokenization and vocabulary handling. GEMORNA-UTR was generated using its native generative procedure with stochastic decoding. All comparator methods used the same predefined target-length vector as EukaUTR.

##### Sequence normalization, nucleotide composition and predicted RNA structure

Before evaluation, sequences were standardized to the RNA alphabet by replacing T with U, and analyses were restricted to the four canonical RNA nucleotides. Unless otherwise specified, the length-matched natural set was used as the reference.

GC content was calculated as the fraction of nucleotides that were G or C. Predicted RNA secondary-structure propensity was evaluated using RNAfold from the ViennaRNA package with the –noPS option. Minimum free energy (MFE) was divided by sequence length and reported in kcal mol^−1^ nt^−1^. More negative values indicate lower predicted folding free energy per nucleotide. RNAfold-derived MFE was treated as a computational structural prediction rather than an experimentally measured property.

##### Masked-language-model naturalness score

Sequence naturalness was quantified using EukaUTR-S2 as a fixed masked-language-model evaluator. Each nucleotide position was masked individually, and the probability assigned to the original nucleotide was recorded. For a sequence **x** = (*x*_1_*, …, x_N_*), the score was defined as

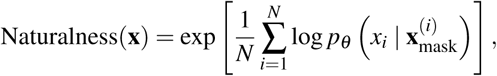

where 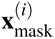 denotes the sequence with nucleotide *i* replaced by <mask>. Padding and special-token positions were excluded. Up to 1,000 sequences were sampled from each method using random seed 42, and masked positions were evaluated in inference batches of 64. Higher scores indicate greater compatibility with the sequence distribution learned by EukaUTR-S2. Because EukaUTR-S2 was itself evaluated as a generator, this score was treated as a model-based measure of sequence compatibility rather than an independent measure of biological function.

##### *k*-mer composition

Local sequence composition was evaluated using nucleotide *k*-mer frequency profiles for *k* = 3–6. Counts were pooled across sequences within each set and normalized to frequencies, with absent *k*-mers assigned a frequency of zero. Similarity to the natural reference was quantified primarily by Pearson correlation between the generated and natural *k*-mer frequency profiles. Cosine similarity, *L*_1_ distance and total-variation distance were calculated as complementary measures, whereas Pearson correlation was used for the primary comparison in Fig. 5.

##### Canonical cis-regulatory motif density

Four classes of canonical 3′ UTR regulatory motifs were evaluated: polyadenylation signals (PASs), AU-rich elements (AREs), cytoplasmic polyadenylation elements (CPEs) and Pumilio response elements (PREs). PASs were defined as AAUAAA or AUUAAA. AREs were identified by an AUUUA core within an AU-rich local sequence context; for each occurrence, the core and up to 10 nt on either side were evaluated, and sites with an A/U fraction of at least 0.70 were retained. CPEs were defined as UUUUAU or UUUUAAU, and PREs were identified using the consensus UGUA[ACGU]AUA. Overlapping occurrences were retained, whereas alternative motif definitions corresponding to the same nucleotide interval were counted once.

For each motif class, density was calculated as the total number of detected occurrences divided by the total evaluated sequence length and reported as occurrences per kilobase. Overall deviation from the natural motif profile was summarized as the mean absolute log_2_ fold difference in density across the four motif classes, using a pseudocount of 10^−6^. Lower values indicate closer agreement with the natural-reference motif profile.

##### Human miRNA 8-mer target-site density

Human mature miRNA sequences were obtained from miRBase release 23 [41]. Nucleotides 2–8 of each mature miRNA were extracted as the 7-nt seed, and duplicate seed sequences were removed. Canonical 8-mer target motifs were defined as the reverse complement of the 7-nt seed followed by an adenosine. Duplicate target motifs were collapsed, and all overlapping exact matches were counted. Target-site density was reported as the number of detected sites per kilobase of evaluated sequence.

##### Homopolymer and low-complexity properties

For each sequence, the maximum contiguous run length was calculated separately for A, C, G and U. The primary low-complexity comparison focused on A-and U-homopolymers. Deviation from the natural reference was summarized as the mean of the absolute differences in mean maximum A-run and U-run lengths between each generated set and the natural set.

The longest homopolymer of any nucleotide was also recorded for each sequence. Complementary analyses quantified the fraction of sequences containing at least one homopolymeric tract of six or more nucleotides and the fraction whose maximum homopolymer length exceeded the 95th percentile of the corresponding distribution in the complete natural pre-training corpus.

##### Embedding-based distributional similarity

Embedding-based analyses used EukaUTR-S2 as a fixed evaluator. Sequence representations were extracted from the final Transformer layer using the <cls> embedding. The same checkpoint and representation-extraction procedure were applied to all sequence sets, yielding 10,000 embeddings per set.

Before distance calculation, each embedding dimension was standardized using the mean and standard deviation estimated from the natural-reference embeddings; no dimensionality reduction was applied. Distributional similarity to the natural reference was quantified using squared maximum mean discrepancy (MMD^2^) with a radial basis function kernel [38] and Fréchet distance (FD) [37].

For MMD^2^, the kernel bandwidth was fixed at *σ* = 10. For each method-to-natural comparison, 1,024 embeddings were sampled from each distribution and MMD^2^ was recalculated over 100 iterations using random seed 42. The mean across iterations was used as the primary estimate, and the empirical standard deviation and 2.5th and 97.5th percentiles were retained as uncertainty summaries.

FD was calculated using all available standardized embeddings. Covariance matrices were symmetrized and regularized using a shrinkage coefficient of 10^−3^ and diagonal jitter of 10^−6^. Matrix square roots were computed by eigendecomposition, with small negative eigenvalues arising from numerical imprecision clipped to zero. Lower MMD^2^ and FD values indicate closer agreement with the natural embedding distribution.

As an internal reference, natural embeddings were divided into independent, non-overlapping subsets and evaluated using the same procedures. Because all embedding-based metrics were calculated in EukaUTR-S2 embedding space, they were interpreted as internal measures of distributional similarity and considered together with the direct sequence-level analyses. The overall distributional-evaluation framework followed Strashnov et al. [42].

##### Nearest-neighbour sequence similarity

Nearest-neighbour sequence similarity was evaluated using MMseqs2 [22] both within each generated set and against the complete EukaUTR pre-training corpus. For within-set searches, self-matches were excluded and the highest-identity non-self match was identified for each sequence. Matches were counted when they reached at least 80% sequence identity and 0.8 alignment coverage. The same thresholds were used when searching generated sequences against the complete pre-training corpus. Within-set frequencies were reported per 10,000 evaluated sequences.

#### EukaUTR-Guide implementation and evaluation

##### Construction of steering and reference sequence sets

EukaUTR-Guide was inspired by steering-vector approaches to protein sequence design [39], but uses the resulting steering direction only to prioritize editing positions rather than to modify model activations during decoding. Candidate sequences were derived from genomic 3′ UTRs associated with the 5,000 most highly expressed genes in Cancer Cell Line Encyclopedia (CCLE) expression profiles [43]. After sequence retrieval and filtering, 3,328 3′ UTRs were retained and divided into overlapping 110-nt fragments with a stride of 30 nt, yielding 38,916 fragments.

Each fragment was scored using an EukaUTR-S2 degradation-rate predictor fine-tuned on the Rabani et al. UTR-seq degradation-rate benchmark [2], yielding a predicted degradation rate *β̂*. Fragments in the lower and upper 5% of the predicted-rate distribution were used as candidate low-and high-rate steering sequences, respectively. From these pools, 100 low-rate fragments were selected to represent the desired low-degradation state and 100 high-rate fragments to represent the contrasting state. Selected fragments were restricted to no more than two per gene and to pairwise sequence identity of at most 80%. The low-and high-rate steering sets were additionally matched for GC content within five percentage points.

Steering fragments were excluded from subsequent reference-set construction. The remaining fragments were stratified by their initial predicted degradation rates into high (*β̂* > 0.4), medium (0.3 ≤ *β̂* ≤ 0.4) and low (*β̂* < 0.3) groups. From each group, 1,000 reference fragments were sampled while maintaining pairwise sequence identity of at most 80%.

##### Low-degradation steering direction and Guided editing

EukaUTR-S2 was used for both representation extraction and masked-nucleotide decoding, with all pretrained parameters held fixed. For a sequence *x* containing *K* nucleotide tokens, its sequence-level representation at Transformer layer *ℓ* was obtained by mean pooling the nucleotide-level hidden representations,

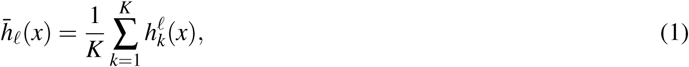

where 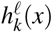 denotes the hidden representation at nucleotide position *k*. Padding and special tokens were excluded from the mean. The low-degradation steering direction was defined as the difference between the mean representations of the low-and high-rate steering sets,

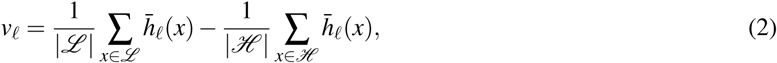

where *L* and *H* denote the low-and high-rate steering sets, respectively. Representations from the final Transformer layer were used throughout.

For each reference sequence, alignment of nucleotide position *k* with the low-degradation steering direction was quantified by cosine similarity,

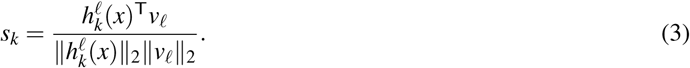

Lower *s_k_* values indicate weaker alignment with the low-degradation direction. In each Guided editing round, the four positions with the lowest scores (*T* = 4) were simultaneously replaced with <mask> and decoded using EukaUTR-S2. Decoding probabilities were restricted to A, C, G and U, and replacement nucleotides were sampled by multinomial sampling at temperature 1.0 without top-*p* truncation.

The steering direction was not added to hidden activations during decoding (activation-steering coefficient *α* = 0). Thus, the steering direction determined where editing was attempted, whereas the replacement nucleotide at each selected position was determined solely by the pretrained masked-language model.

Editing was repeated for eight rounds (*R* = 8). After each round, the edited sequence was re-encoded and position scores were recalculated. Previously selected positions remained eligible for selection in subsequent rounds. Because the masked-language model could sample the original nucleotide, selection of a position did not necessarily result in a nucleotide substitution.

##### Unguided editing control

Unguided editing provided a matched control for steering-direction-based site selection. In each of eight rounds, four valid nucleotide positions were selected at random and decoded using the same EukaUTR-S2 checkpoint, nucleotide restrictions, sampling temperature and decoding procedure as in Guided editing. Guided and Unguided editing therefore differed only in the rule used to select editing positions.

##### Evaluation of editing outcomes and cross-predictor analysis

Reference, Unguided and Guided sequences were scored using the EukaUTR-S2 degradation-rate predictor. Predicted degradation-rate reduction was calculated as the reference rate minus the edited-sequence rate, such that positive values indicate lower predicted degradation after editing. The Guided-versus-Unguided effect was calculated as the Unguided rate minus the Guided rate, with positive values indicating lower predicted degradation after Guided editing. All paired comparisons were performed between sequences derived from the same starting reference sequence. Guided and Unguided outcomes were compared using two-sided paired Wilcoxon signed-rank tests. Holm correction was applied across the three degradation-rate strata for the primary EukaUTR-S2 analysis and separately across the nine external predictor–stratum comparisons for the cross-predictor analysis.Confidence intervals were estimated from 4,000 bootstrap resamples at the starting-reference-sequence level, with paired outcomes resampled together. The median was used as the summary statistic, and the 2.5th and 97.5th percentiles were used as the 95% confidence limits.

Because the EukaUTR-S2 degradation-rate predictor was used both to construct the steering sets and to evaluate the primary editing outcome, the same sequences were additionally evaluated using degradation-rate predictors based on GEMORNA-UTR, mRNABERT and 3UTRBERT. Each external predictor was fine-tuned on the same Rabani et al. degradation-rate benchmark using the same downstream fine-tuning protocol.

Predicted degradation rates were used only for steering-set construction and final outcome evaluation. No degradation-rate predictor was used during iterative editing to accept, reject or rank candidate nucleotide substitutions. EukaUTR-Guide therefore does not rely on predictor-in-the-loop rejection sampling or predictor-based candidate selection.

##### Editing-site and sequence-property analyses

Unless otherwise specified, analyses were performed on the high-rate reference group. A realized substitution was defined as a position at which the final edited sequence differed from its reference. Degradation-rate reduction per substitution was calculated from the reference-minus-edited prediction divided by the number of realized substitutions.

Selected-site analyses used unique nucleotide positions across editing rounds. Reference-to-final nucleotide transitions and nucleotide enrichment were compared between Guided and Unguided editing. Sequence-level changes in GC content, RNAfold-predicted minimum free energy, nearest-neighbour normalized Hamming distance and predicted human miRNA 8-mer target-site abundance were evaluated relative to the corresponding reference sequences. Confi-dence intervals for nucleotide enrichment were obtained from 4,000 bootstrap resamples of reference sequences, with enrichment recalculated for each resample and percentile-based 95% confidence limits reported.

##### GC-content sensitivity analysis

To assess whether GC enrichment contributed to the predicted Guided effect, Guided-versus-Unguided differences in degradation-rate reduction were examined using nested linear models with sequential adjustment for ΔGC and initial predicted degradation rate. Cluster-robust standard errors were calculated at the reference-sequence level. Associations between ΔGC and degradation-rate reduction were assessed using Spearman correlation.

As a complementary analysis, Guided and Unguided sequences were matched one-to-one on ΔGC and initial predicted degradation rate. Covariate balance was assessed using standardized mean differences, and matched outcomes were compared using a two-sided paired Wilcoxon signed-rank test.

### Supplementary Figures

**Supplementary Figure 1:**
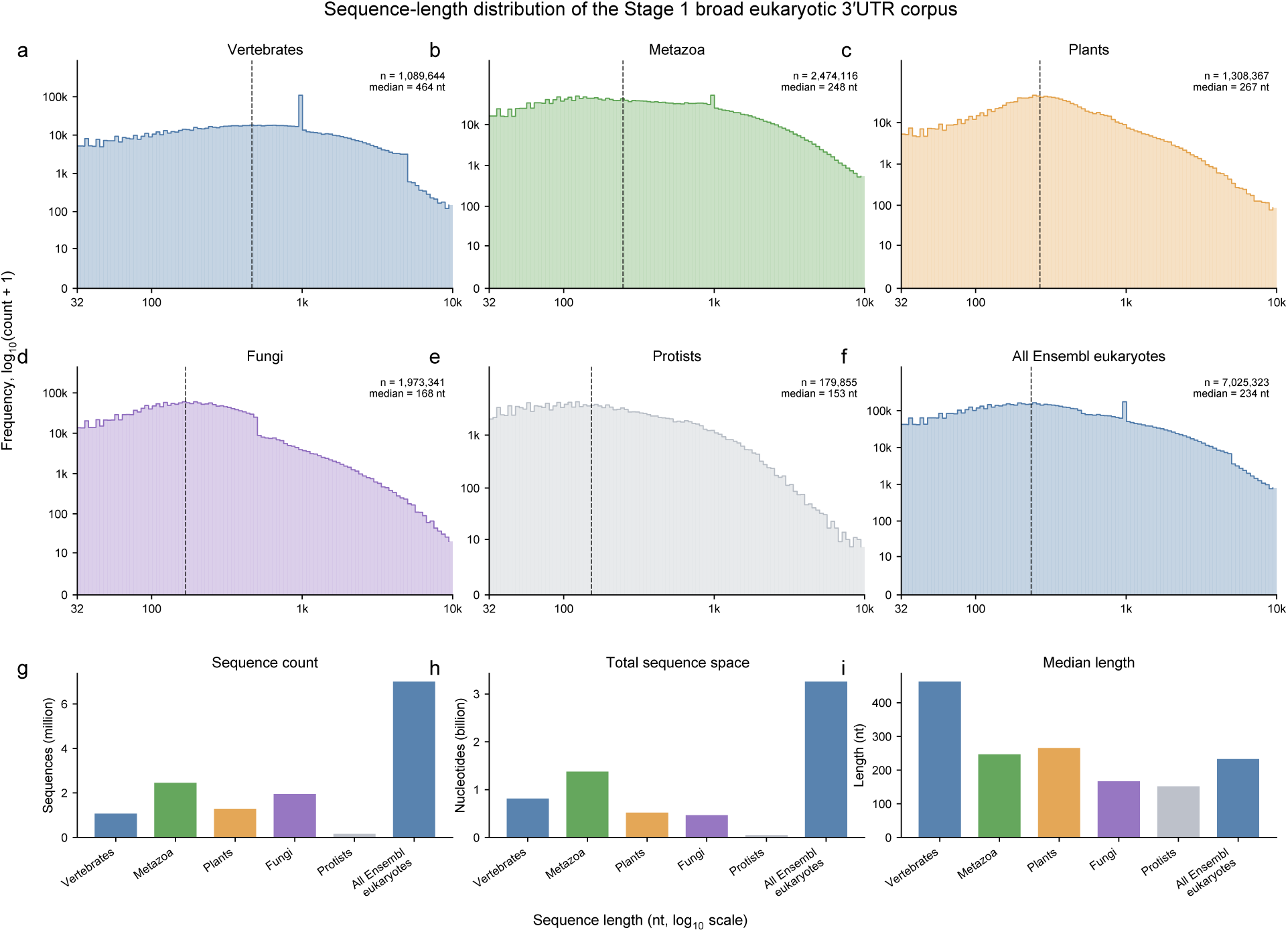
Sequence-length distributions and composition of the Stage 1 broad eukaryotic 3′ UTR corpus. **a–e,** Sequence-length distributions of non-redundant 3′ UTRs from vertebrates (**a**), non-vertebrate metazoans (**b**), plants (**c**), fungi (**d**) and protists (**e**). **f,** Sequence-length distribution of the complete Stage 1 corpus. **g–i,** Number of sequences (**g**), total nucleotide content (**h**) and median sequence length (**i**) for each taxonomic group and the complete corpus. Sequence length and frequency are shown on logarithmic scales in **a–f**. Dashed vertical lines indicate median sequence lengths; sample sizes and medians are shown in the corresponding panels.

**Supplementary Figure 2:**
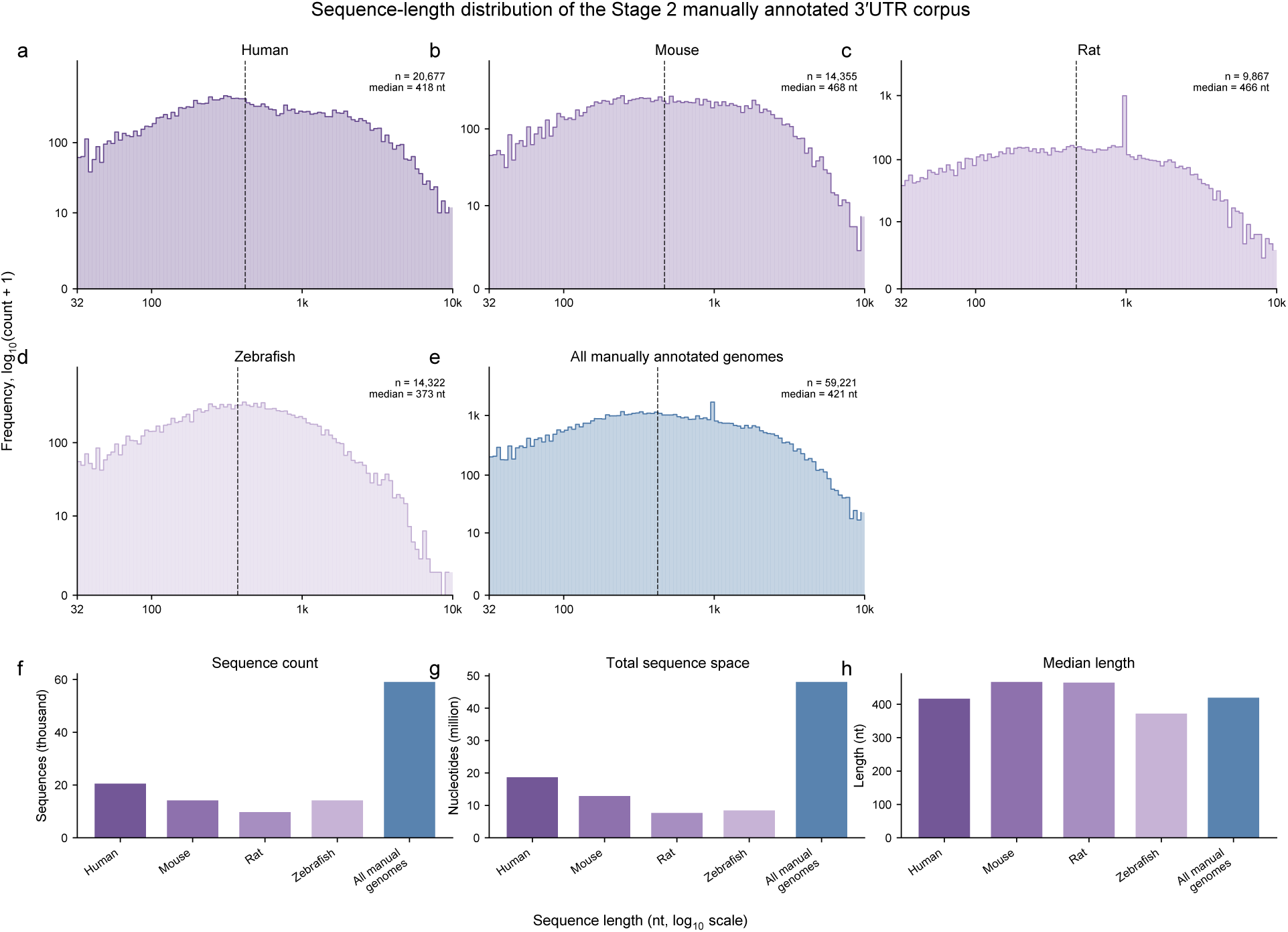
Sequence-length distributions and composition of the Stage 2 high-confidence 3′ UTR corpus. **a–d,** Sequence-length distributions of non-redundant 3′ UTRs from human (**a**), mouse (**b**), rat (**c**) and zebrafish (**d**). **e,** Sequence-length distribution of the complete Stage 2 corpus. **f–h,** Number of sequences (**f**), total nucleotide content (**g**) and median sequence length (**h**) for each species and the complete corpus. Sequence length and frequency are shown on logarithmic scales in **a–e**. Dashed vertical lines indicate median sequence lengths; sample sizes and medians are shown in the corresponding panels.

**Supplementary Figure 3:**
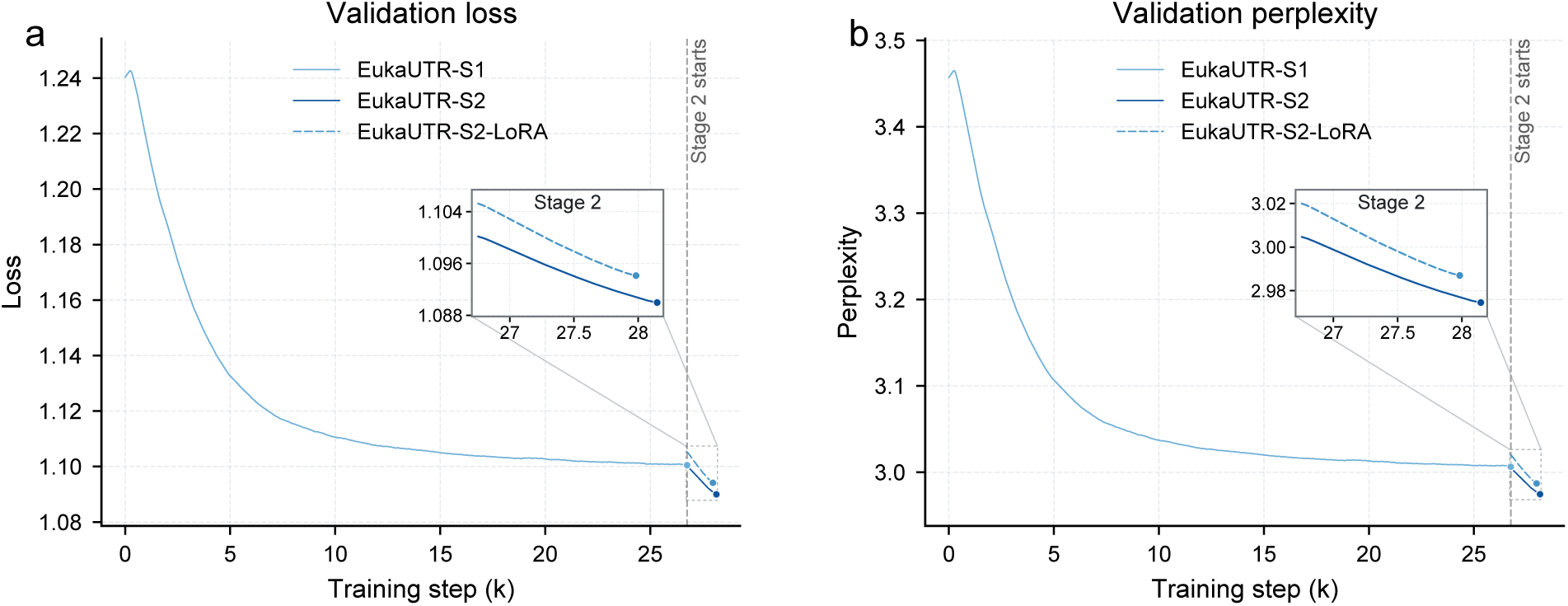
Validation trajectories during EukaUTR pre-training and Stage 2 refinement. **a,b,** Validation loss (**a**) and perplexity (**b**) during Stage 1 pre-training and subsequent Stage 2 refinement by full-parameter continued pre-training (EukaUTR-S2) or LoRA-based adaptation (EukaUTR-S2-LoRA). The vertical dashed line marks the transition from Stage 1 to Stage 2, and the insets show the Stage 2 trajectories at higher resolution. Training steps are shown in thousands.

**Supplementary Figure 4:**
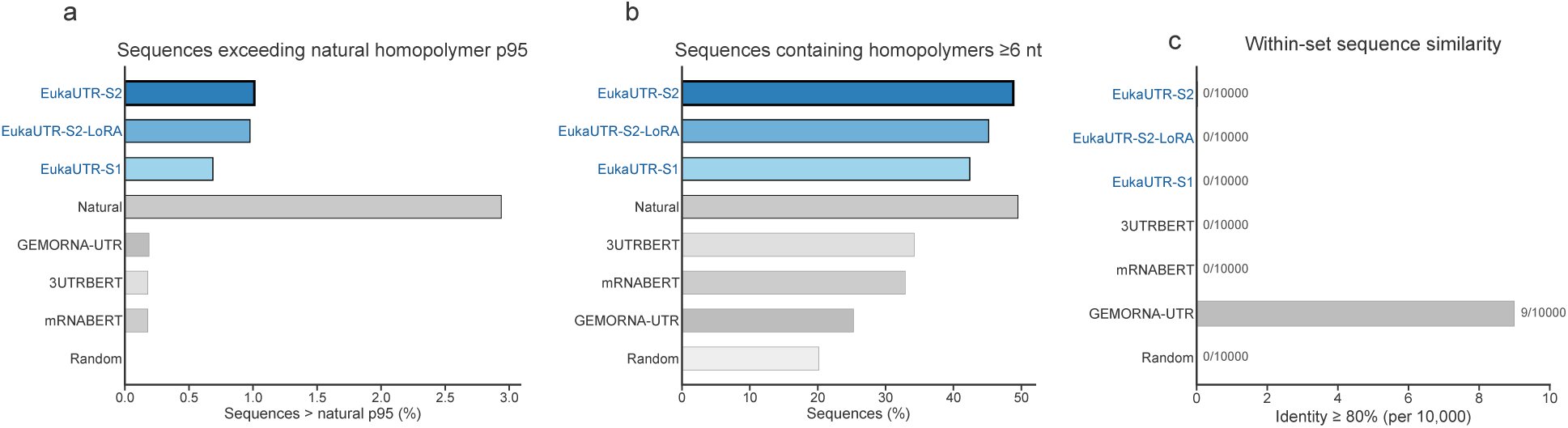
Homopolymer properties and within-set sequence similarity of de novo-generated 3′ UTRs. **a,** Percentage of sequences whose maximum homopolymer length exceeded the 95th percentile of the maximum-homopolymer-length distribution in the complete natural pre-training corpus. **b,** Percentage of sequences containing at least one homopolymeric tract of ≥ 6 nt. **c,** Within-set nearest-neighbour sequence similarity assessed using MMseqs2. For each generated sequence, the highest-identity non-self match within the same generated set was identified using a minimum alignment coverage of 0.8. Bars show the frequency of sequences whose nearest-neighbour match reached ≥ 80% sequence identity, reported per 10,000 evaluated sequences.

**Supplementary Figure 5:**
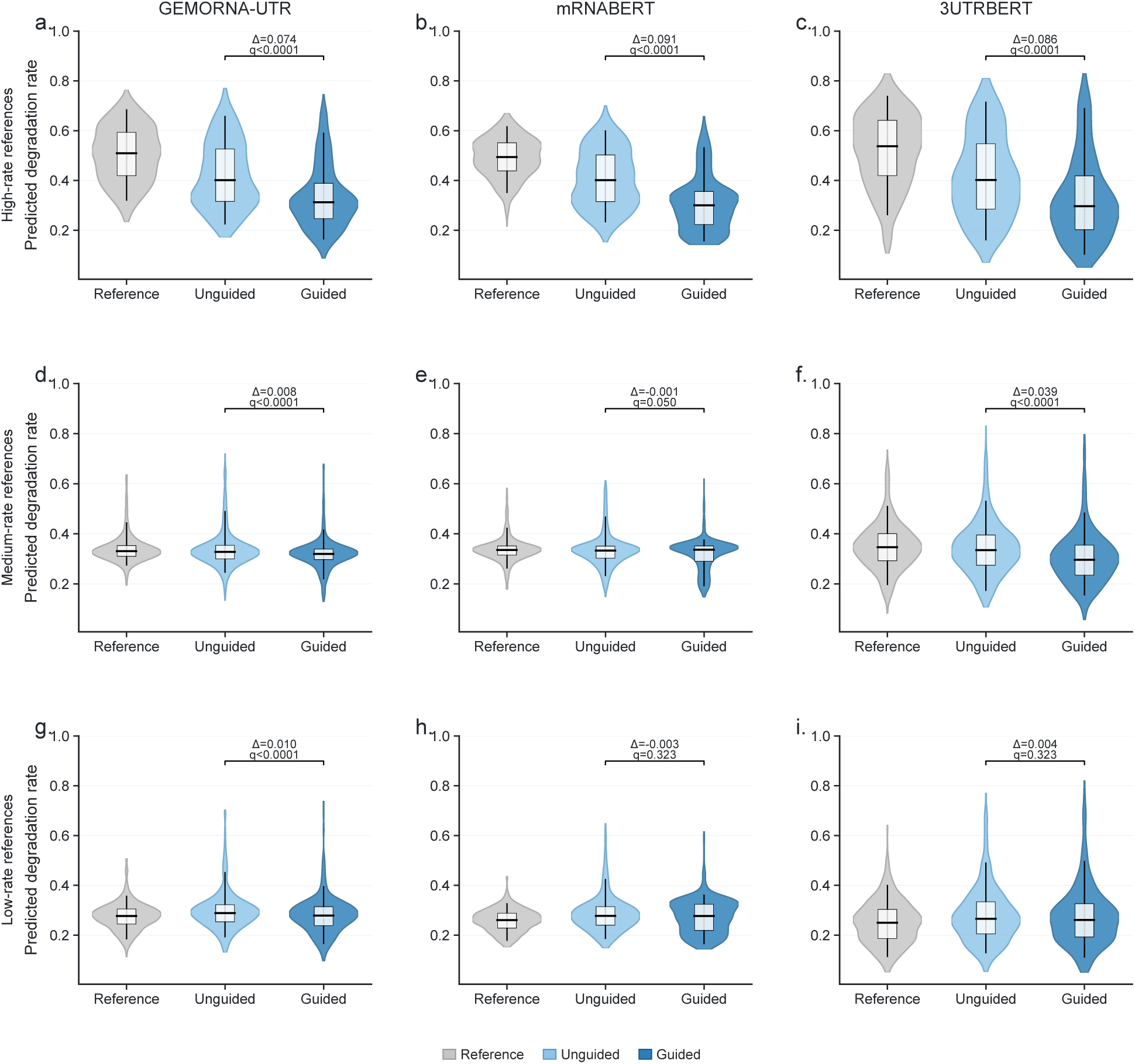
Cross-predictor evaluation of EukaUTR-Guide editing. Predicted degradation rates for Reference, Unguided and Guided sequences were independently evaluated using degradation-rate predictors based on GEMORNA-UTR (**a,d,g**), mRNABERT (**b,e,h**) and 3UTRBERT (**c,f,i**). Rows correspond to reference sequences with high (**a–c**), medium (**d–f**) and low (**g–i**) initial predicted degradation rates. Within each stratum, Reference, Unguided and Guided sequences are paired by their common starting reference sequence. Δ denotes the median paired difference *β̂*_Unguided_ − *β̂*_Guided_; positive values indicate lower predicted degradation after Guided editing. Adjusted *P* values are from two-sided paired Wilcoxon signed-rank tests with Holm correction across the nine predictor–stratum comparisons. Violins show distributions, internal boxes indicate medians and interquartile ranges, and whiskers extend from the 5th to the 95th percentiles.

**Supplementary Figure 6:**
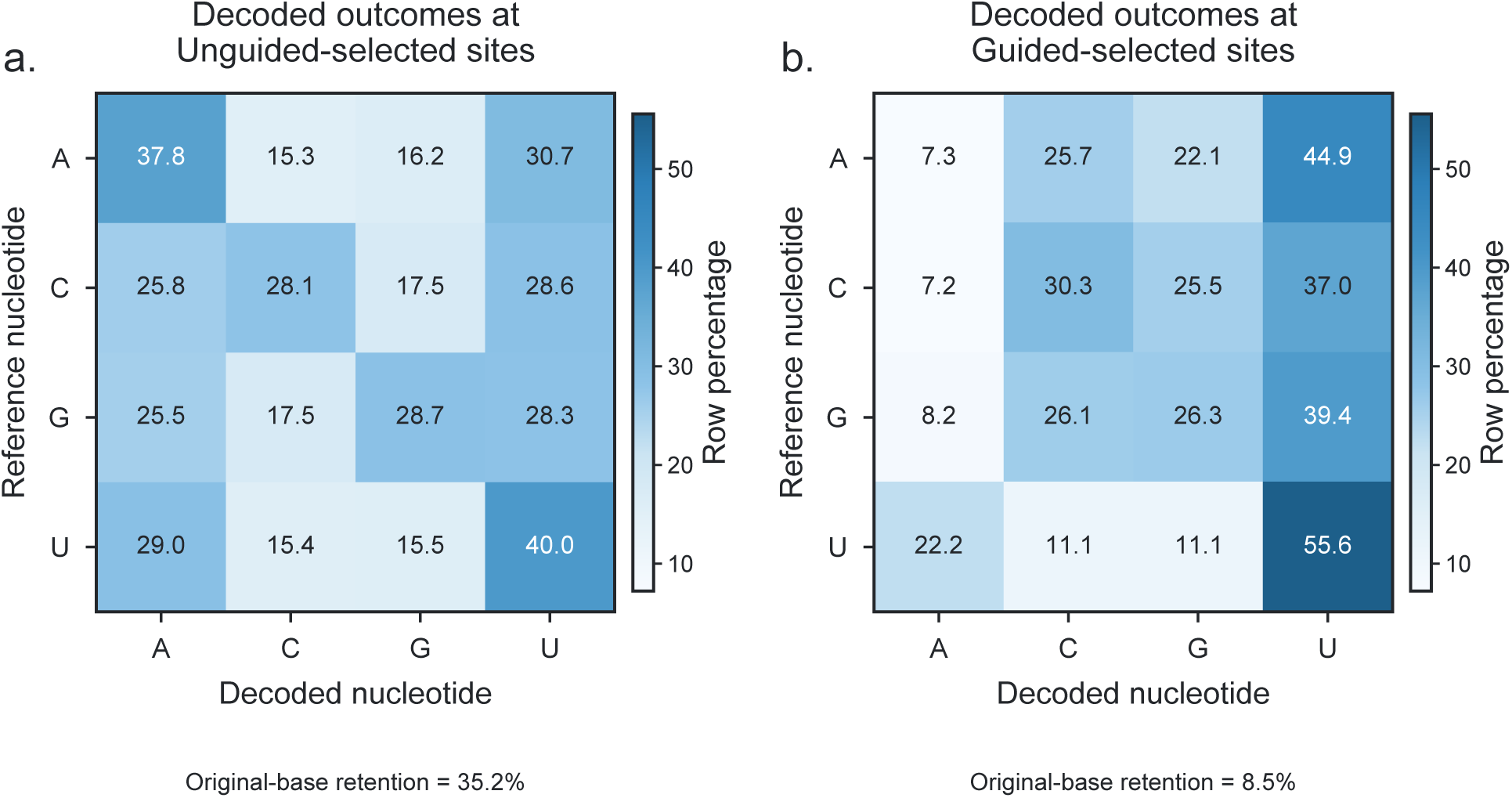
Nucleotide outcomes at sites selected during EukaUTR-Guide editing. Analyses were performed on the high-rate reference group. **a,b,** Reference-to-final nucleotide transitions at unique positions selected during Unguided (**a**) and Guided (**b**) editing. Values are row-normalized percentages, and diagonal entries indicate retention of the reference nucleotide. The original nucleotide was retained at 35.2% of Unguided-selected positions and 8.5% of Guided-selected positions.

**Supplementary Figure 7:**
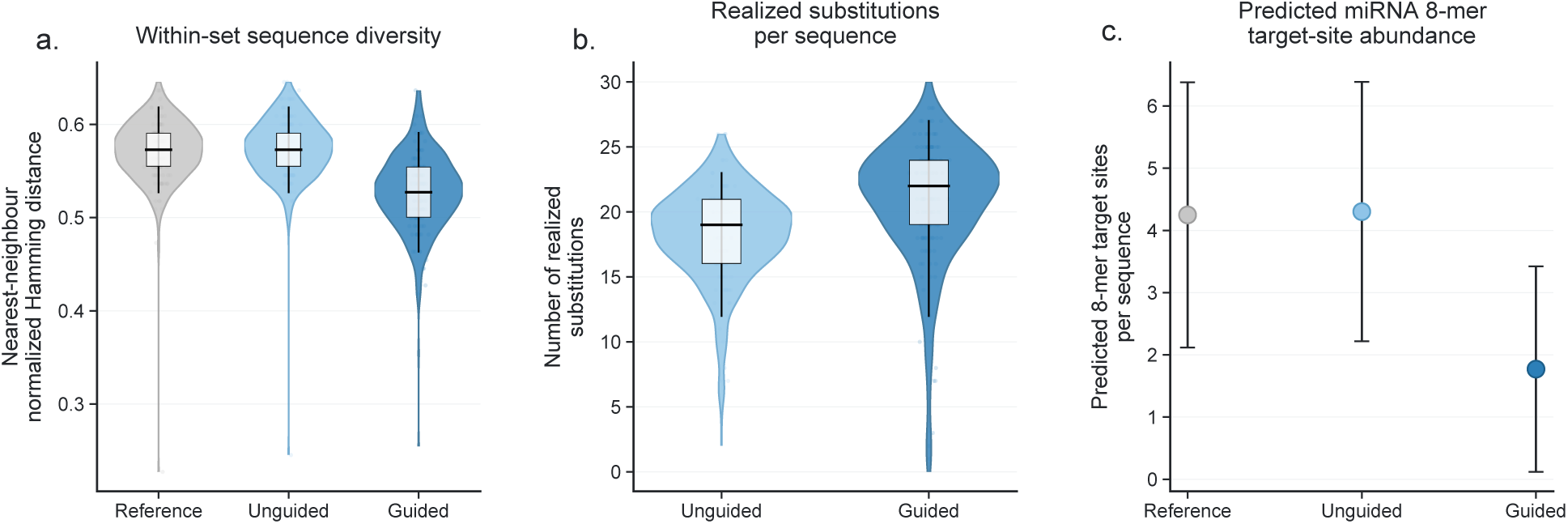
Additional sequence properties following EukaUTR-Guide editing. Analyses were performed on the high-rate reference group. **a,** Within-set sequence diversity of Reference, Unguided and Guided sequences, quantified by nearest-neighbour normalized Hamming distance. **b,** Number of realized nucleotide substitutions per sequence after Unguided or Guided editing. **c,** Predicted human miRNA 8-mer target-site abundance in Reference, Unguided and Guided sequences. Points indicate group means and error bars denote s.d. In **a,b**, violins show distributions, internal boxes indicate medians and interquartile ranges, and whiskers extend from the 5th to the 95th percentiles.

**Supplementary Figure 8:**
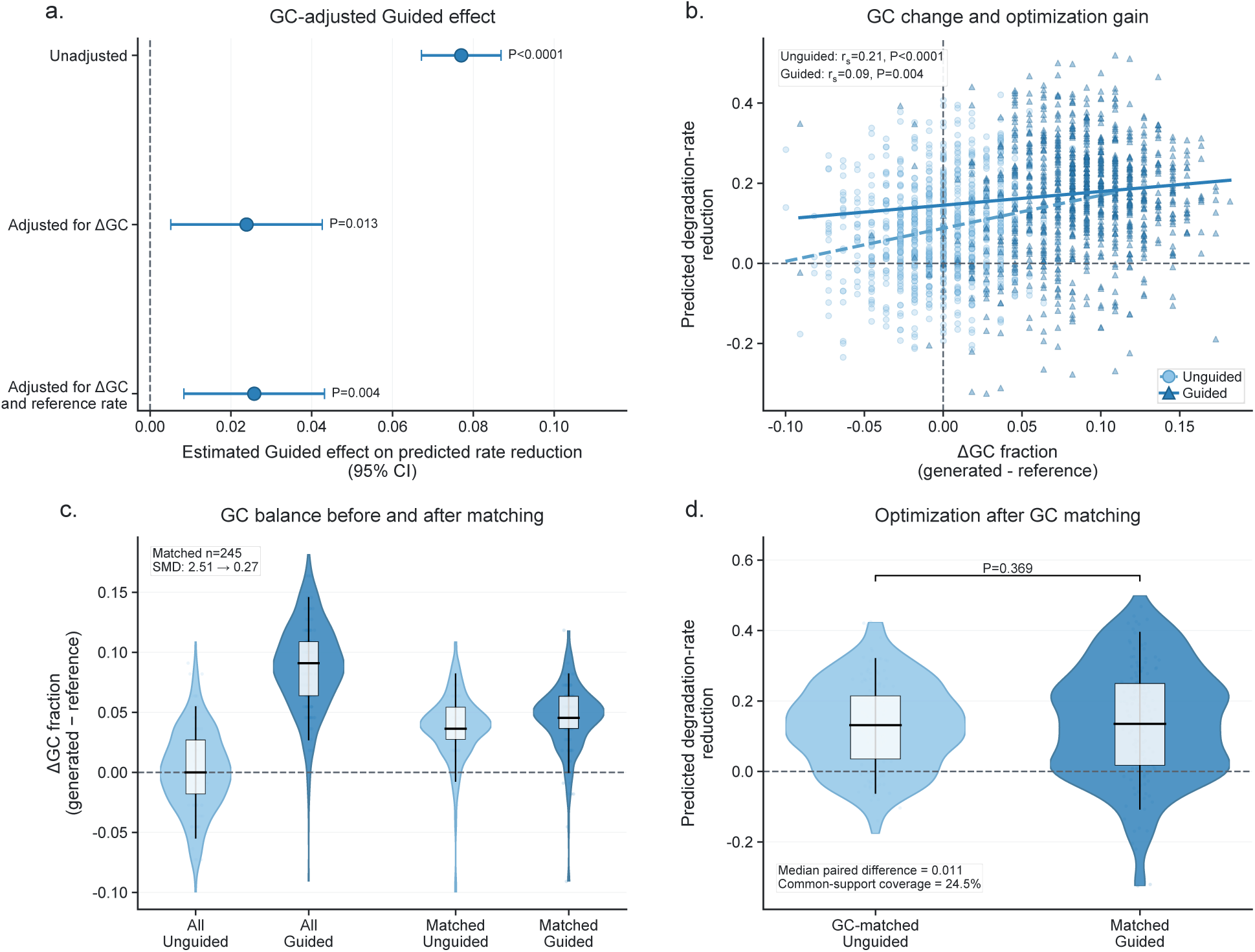
GC-content sensitivity analysis of the predicted EukaUTR-Guide editing effect. Analyses were performed on the high-rate reference group. **a,** Estimated Guided-versus-Unguided difference in predicted degradation-rate reduction from nested regression models including editing method alone, additionally adjusting for ΔGC, or additionally adjusting for both ΔGC and the initial predicted degradation rate. Points indicate regression coefficients for Guided versus Unguided editing, and error bars denote 95% confidence intervals calculated using cluster-robust standard errors at the reference-sequence level. **b,** Association between ΔGC and predicted degradation-rate reduction for Unguided and Guided editing. Spearman correlation coefficients and corresponding *P* values are shown. **c,** Distributions of ΔGC before and after one-to-one matching of Unguided and Guided sequences on ΔGC and initial predicted degradation rate. SMD denotes standardized mean difference. **d,** Predicted degradation-rate reduction in the 245 matched Unguided–Guided pairs. The displayed *P* value is from a two-sided paired Wilcoxon signed-rank test. Common-support coverage denotes the proportion of Guided sequences retained after matching. In **c,d**, violins show distributions, internal boxes indicate medians and interquartile ranges, and whiskers extend from the 5th to the 95th percentiles.

### Supplementary Tables

**Supplementary Table 1.** Composition of the Stage 1 broad eukaryotic 3′ UTR pre-training corpus.

| Taxonomic group | Genomes | 3' UTRs | Mean length (nt) | Median length (nt) | Total nt (10<sup>9</sup>) |
| --- | --- | --- | --- | --- | --- |
| Vertebrates | 319 | 1,089,644 | 753.21 | 464 | 0.821 |
| Non-vertebrate metazoans | 353 | 2,474,116 | 560.40 | 248 | 1.386 |
| Plants | 198 | 1,308,367 | 403.77 | 267 | 0.528 |
| Fungi | 719 | 1,973,341 | 241.24 | 168 | 0.476 |
| Protists | 126 | 179,855 | 315.79 | 153 | 0.057 |
| **Total** | **1,715** | **7,025,323** | **465.22** | **234** | **3.268** |

Statistics are reported after removing sequences shorter than 32 nt or longer than 10,000 nt and reducing redundancy using MMseqs2 at 80% sequence identity, with one representative retained per cluster. Human, mouse, rat and zebrafish were excluded from Stage 1 and reserved for Stage 2.

**Supplementary Table 2.**
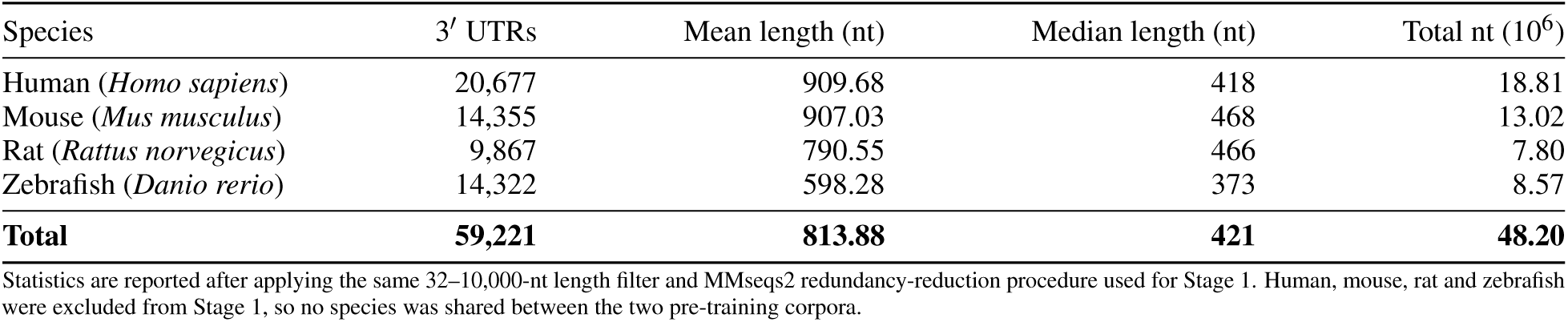
Composition of the Stage 2 high-confidence 3′ UTR pre-training corpus.

**Supplementary Table 3.**
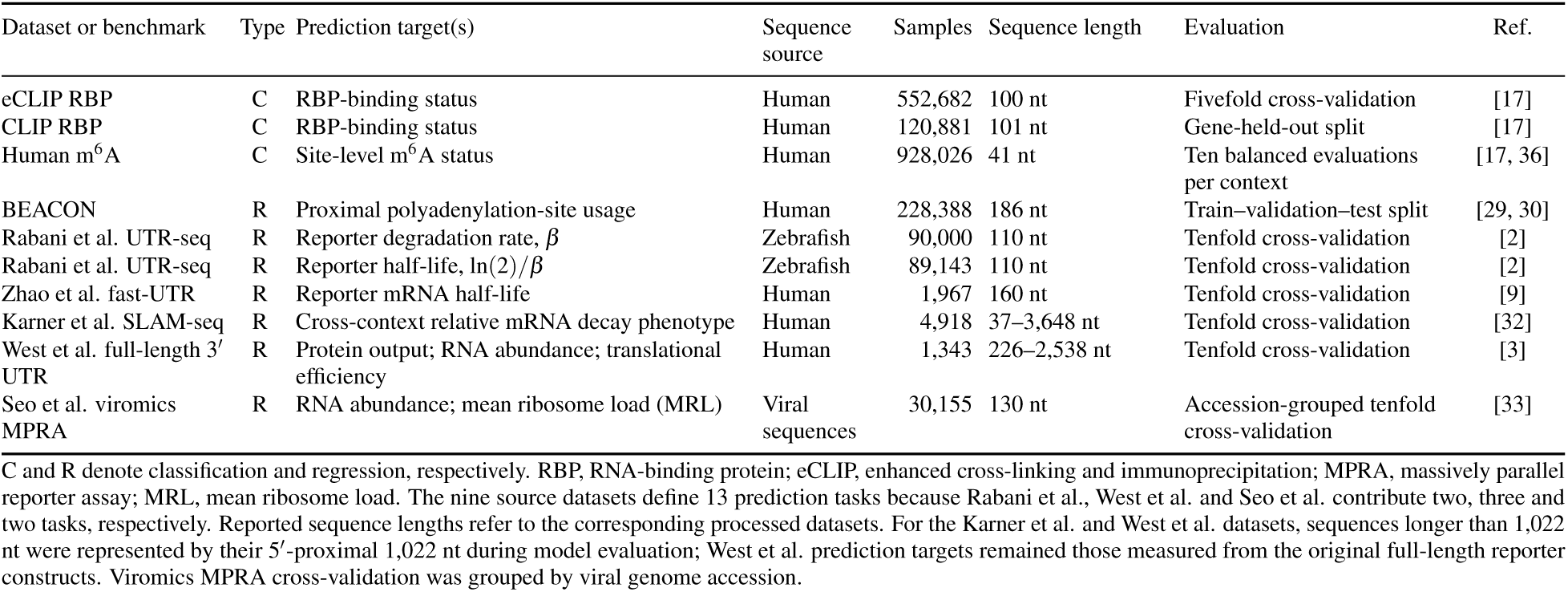
Summary of downstream datasets and prediction tasks.

**Supplementary Table 4.**
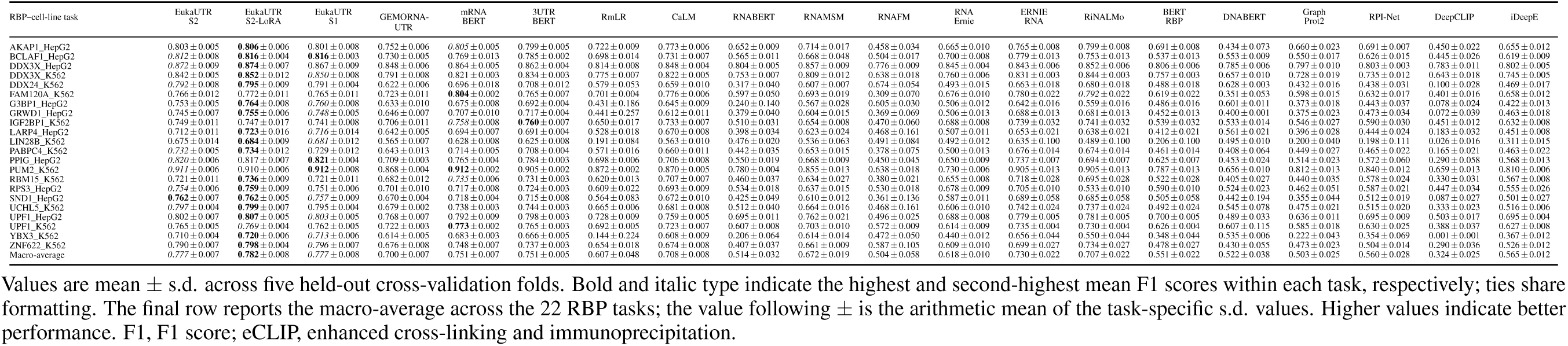
F1 scores for RBP-binding prediction acros 22 eCLIP RBP tasks.

**Supplementary Table 5.**
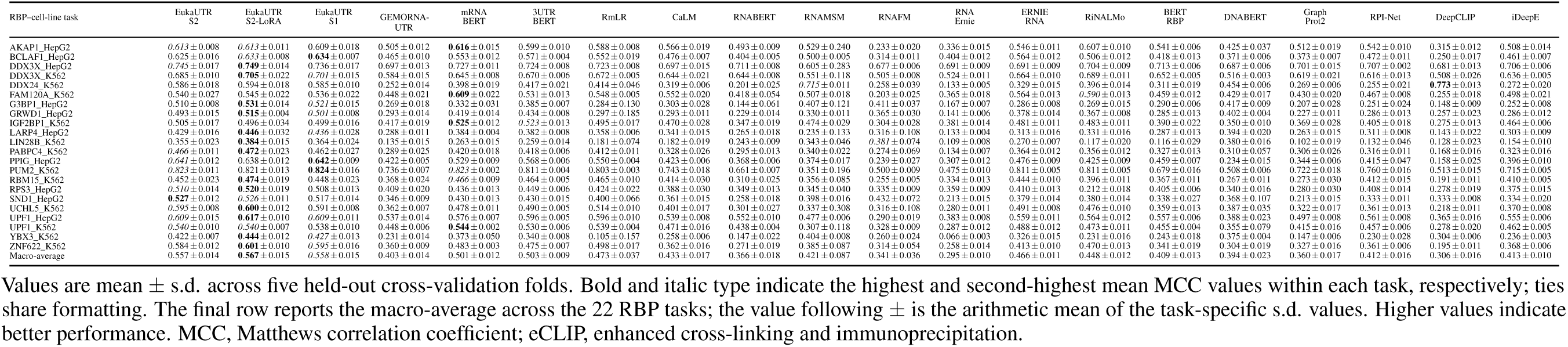
Matthews correlation coefficients for RBP-binding prediction across 22 eCLIP RBP tasks.

**Supplementary Table 6.**
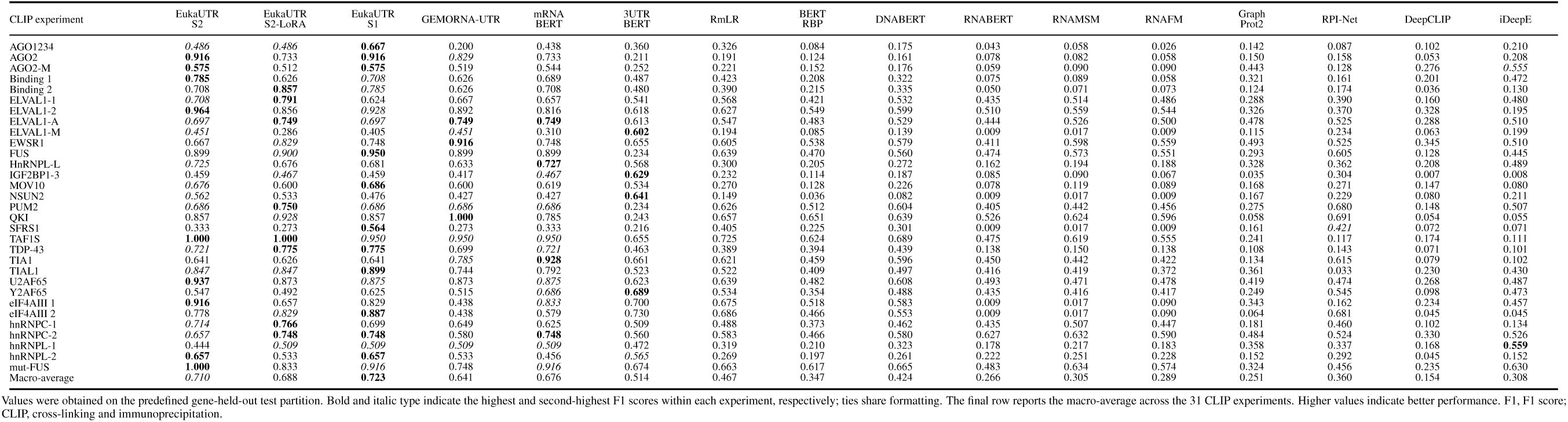
F1 scores for RBP-binding prediction across 31 gene-held-out CLIP experiments.

**Supplementary Table 7.**
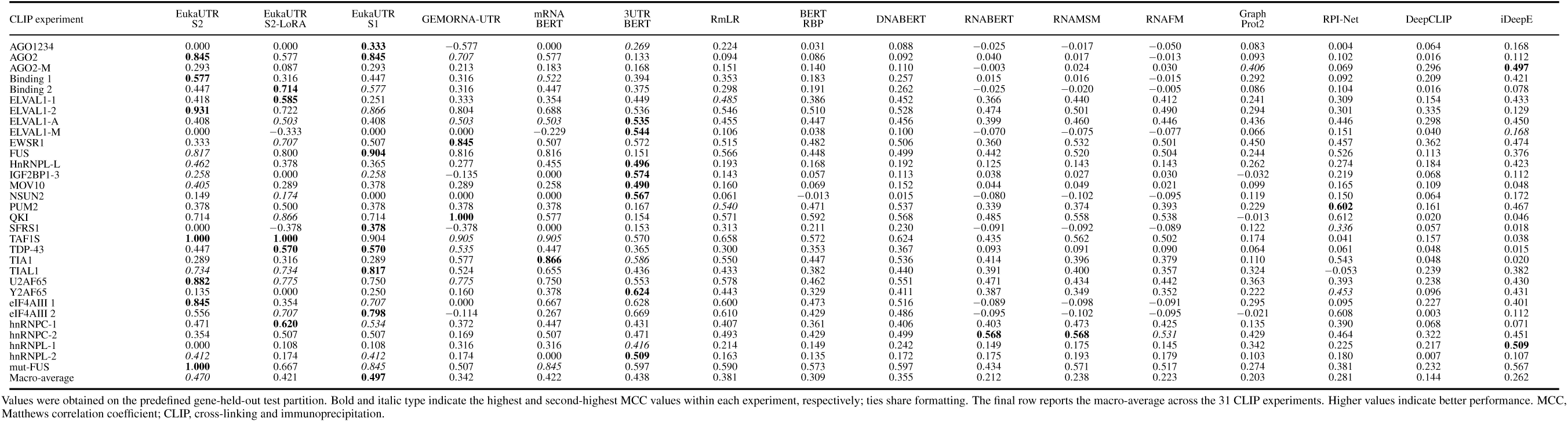
Matthews correlation coefficients for RBP-binding prediction across 31 gene-held-out CLIP experiments.

**Supplementary Table 8.** F1 scores for m^6^A-site prediction across nine human cell lines.

| Cellular context | EukaUTR-S2 | EukaUTR-S2-LoRA | EukaUTR-S1 | GEMORNA-UTR | mRNABERT | 3UTRBERT | DeepM6ASeq | WHISTLE | iMRM | SCRAMP |
| --- | --- | --- | --- | --- | --- | --- | --- | --- | --- | --- |
| A549 | 0.961 ± 0.005 | 0.954 ± 0.007 | 0.956 ± 0.007 | 0.955 ± 0.005 | **0.964** ± 0.003 | 0.963 ± 0.004 | 0.905 ± 0.008 | 0.860 ± 0.002 | 0.863 ± 0.003 | 0.851 ± 0.005 |
| CD8T | **0.967** ± 0.003 | 0.964 ± 0.004 | 0.963 ± 0.003 | 0.963 ± 0.003 | 0.965 ± 0.003 | 0.965 ± 0.003 | 0.947 ± 0.003 | 0.838 ± 0.001 | 0.839 ± 0.006 | 0.580 ± 0.023 |
| ESC | **0.992** ± 0.004 | 0.990 ± 0.003 | 0.990 ± 0.004 | 0.991 ± 0.003 | 0.989 ± 0.004 | 0.991 ± 0.002 | 0.960 ± 0.005 | 0.237 ± 0.004 | 0.790 ± 0.026 | 0.626 ± 0.017 |
| HCT116 | 0.962 ± 0.005 | 0.961 ± 0.006 | 0.961 ± 0.007 | 0.951 ± 0.004 | 0.963 ± 0.006 | **0.969** ± 0.004 | 0.907 ± 0.008 | 0.703 ± 0.002 | 0.821 ± 0.009 | 0.478 ± 0.054 |
| HEK293 | **0.967** ± 0.004 | 0.963 ± 0.004 | 0.963 ± 0.004 | 0.962 ± 0.004 | 0.961 ± 0.001 | 0.965 ± 0.002 | 0.941 ± 0.006 | 0.827 ± 0.001 | 0.848 ± 0.005 | 0.806 ± 0.006 |
| HEK293T | 0.966 ± 0.002 | 0.964 ± 0.001 | 0.964 ± 0.002 | 0.964 ± 0.002 | **0.967** ± 0.001 | 0.964 ± 0.001 | 0.956 ± 0.002 | 0.572 ± 0.001 | 0.808 ± 0.004 | 0.404 ± 0.016 |
| HeLa | **0.967** ± 0.004 | 0.965 ± 0.003 | 0.964 ± 0.003 | 0.964 ± 0.004 | 0.964 ± 0.004 | 0.966 ± 0.003 | 0.946 ± 0.003 | 0.744 ± 0.002 | 0.832 ± 0.004 | 0.540 ± 0.024 |
| HepG2 | **0.967** ± 0.002 | 0.963 ± 0.002 | 0.964 ± 0.003 | 0.963 ± 0.003 | 0.961 ± 0.002 | 0.965 ± 0.002 | 0.945 ± 0.002 | 0.438 ± 0.001 | 0.817 ± 0.006 | 0.348 ± 0.076 |
| MOLM13 | **0.966** ± 0.002 | 0.965 ± 0.002 | 0.964 ± 0.002 | 0.964 ± 0.002 | 0.963 ± 0.001 | 0.965 ± 0.001 | 0.945 ± 0.002 | 0.910 ± 0.001 | 0.882 ± 0.004 | 0.873 ± 0.003 |
| Macro-average | **0.968** ± 0.003 | 0.965 ± 0.004 | 0.966 ± 0.004 | 0.964 ± 0.003 | 0.966 ± 0.003 | **0.968** ± 0.002 | 0.939 ± 0.004 | 0.681 ± 0.002 | 0.833 ± 0.007 | 0.612 ± 0.025 |

Values are mean ± s.d. across ten approximately balanced positive–negative evaluations within each cellular context. Bold and italic type indicate the highest and second-highest context-specific mean F1 scores, respectively; ties share formatting. The final row reports the macro-average across the nine cell lines; the value following ± is the arithmetic mean of the context-specific s.d. values. Higher values indicate better performance.

**Supplementary Table 9.** Matthews correlation coefficients for m^6^A-site prediction across nine human cell lines.

| Cell lines | EukaUTR-S2 | EukaUTR-S2-LoRA | EukaUTR-S1 | GEMORNA-UTR | mRNABERT | 3UTRBERT | DeepM6ASeq | WHISTLE | iMRM | SCRAMP |
| --- | --- | --- | --- | --- | --- | --- | --- | --- | --- | --- |
| A549 | 0.922 ± 0.011 | 0.910 ± 0.014 | 0.913 ± 0.013 | 0.912 ± 0.010 | **0.929** ± 0.005 | 0.928 ± 0.007 | 0.849 ± 0.017 | 0.776 ± 0.005 | 0.751 ± 0.008 | 0.728 ± 0.009 |
| CD8T | **0.934** ± 0.006 | 0.929 ± 0.009 | 0.928 ± 0.007 | 0.929 ± 0.005 | 0.931 ± 0.005 | 0.931 ± 0.005 | 0.913 ± 0.005 | 0.747 ± 0.002 | 0.712 ± 0.010 | 0.316 ± 0.014 |
| ESC | **0.985** ± 0.008 | 0.980 ± 0.005 | 0.980 ± 0.008 | 0.981 ± 0.005 | 0.981 ± 0.006 | 0.983 ± 0.004 | 0.941 ± 0.009 | 0.257 ± 0.008 | 0.639 ± 0.016 | 0.172 ± 0.022 |
| HCT116 | 0.926 ± 0.009 | 0.923 ± 0.012 | 0.923 ± 0.014 | 0.903 ± 0.008 | 0.928 ± 0.007 | **0.939** ± 0.008 | 0.833 ± 0.017 | 0.602 ± 0.006 | 0.692 ± 0.017 | 0.169 ± 0.024 |
| HEK293 | **0.933** ± 0.007 | 0.928 ± 0.008 | 0.928 ± 0.007 | 0.926 ± 0.007 | 0.922 ± 0.006 | 0.932 ± 0.004 | 0.911 ± 0.013 | 0.735 ± 0.003 | 0.725 ± 0.008 | 0.648 ± 0.008 |
| HEK293T | 0.932 ± 0.003 | 0.931 ± 0.003 | 0.931 ± 0.003 | 0.930 ± 0.003 | **0.937** ± 0.001 | 0.929 ± 0.001 | 0.912 ± 0.003 | 0.490 ± 0.004 | 0.668 ± 0.007 | 0.126 ± 0.007 |
| HeLa | **0.934** ± 0.007 | 0.932 ± 0.006 | 0.931 ± 0.006 | 0.930 ± 0.007 | 0.930 ± 0.004 | 0.933 ± 0.005 | 0.913 ± 0.006 | 0.642 ± 0.005 | 0.701 ± 0.007 | 0.183 ± 0.015 |
| HepG2 | 0.931 ± 0.004 | 0.929 ± 0.003 | 0.930 ± 0.005 | 0.929 ± 0.005 | 0.924 ± 0.003 | **0.932** ± 0.004 | 0.911 ± 0.004 | 0.388 ± 0.007 | 0.684 ± 0.011 | 0.075 ± 0.010 |
| MOLM13 | **0.934** ± 0.003 | 0.931 ± 0.004 | 0.930 ± 0.004 | 0.930 ± 0.004 | 0.925 ± 0.005 | 0.931 ± 0.003 | 0.909 ± 0.005 | 0.843 ± 0.002 | 0.776 ± 0.007 | 0.757 ± 0.007 |
| Macro-average | 0.937 ± 0.006 | 0.933 ± 0.007 | 0.933 ± 0.007 | 0.930 ± 0.006 | 0.934 ± 0.005 | **0.938** ± 0.004 | 0.899 ± 0.009 | 0.609 ± 0.005 | 0.705 ± 0.010 | 0.353 ± 0.013 |

Values are mean ± s.d. across ten approximately balanced positive–negative evaluations within each cell lines. Bold and italic type indicate the highest and second-highest context-specific mean MCC values, respectively; ties share formatting. The final row reports the macro-average across the nine cell lines; the value following ± is the arithmetic mean of the context-specific s.d. values. Higher values indicate better performance. MCC, Matthews correlation coefficient.

**Supplementary Table 10.**
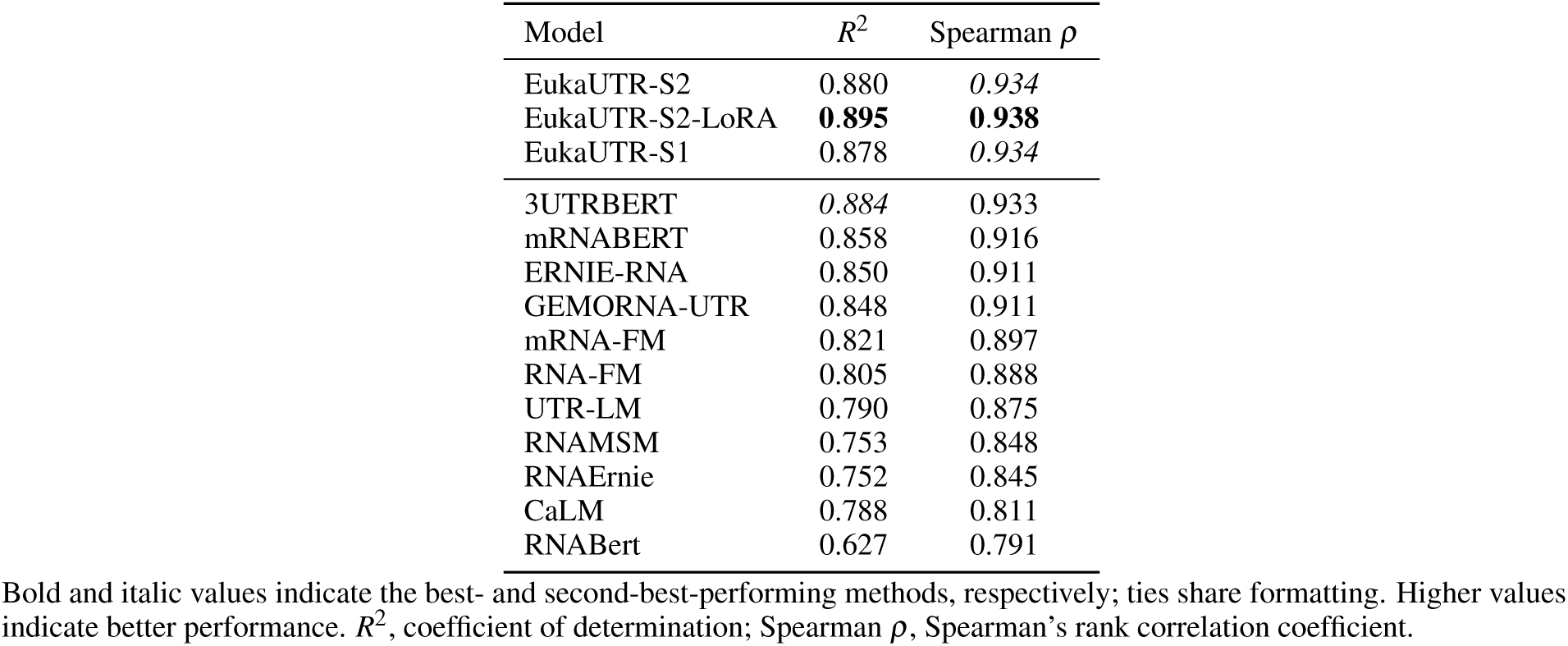
Performance in predicting proximal polyadenylation-site usage on the BEACON benchmark [30].

**Supplementary Table 11.**
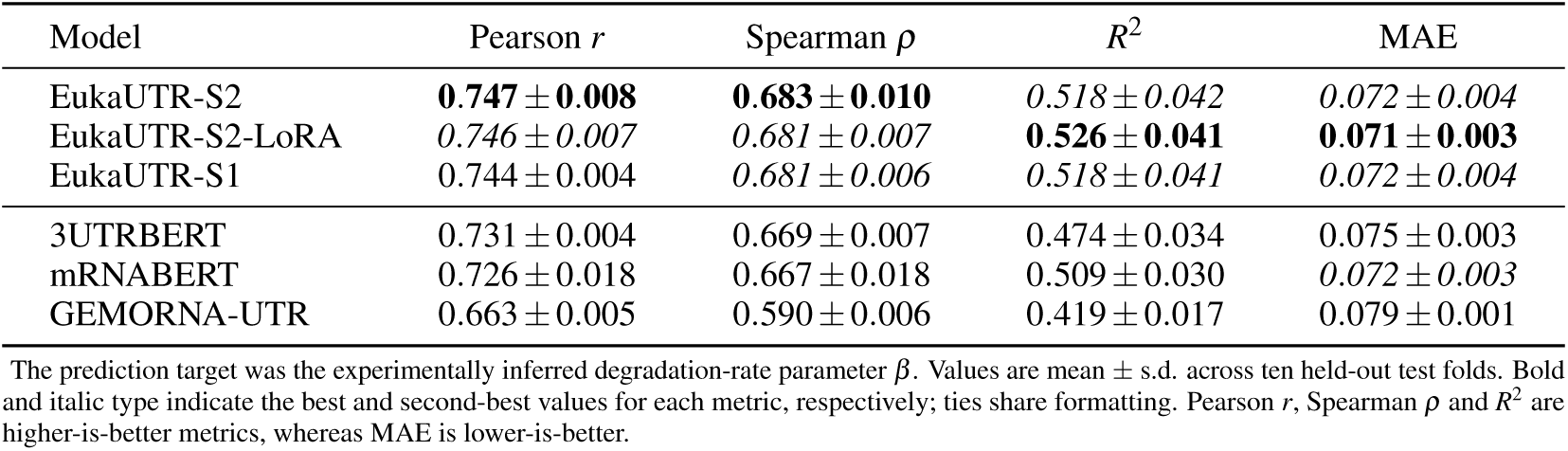
Performance on zebrafish reporter mRNA degradation-rate prediction using the Rabani et al. UTR-seq benchmark.

**Supplementary Table 12.** Performance on derived zebrafish reporter mRNA half-life prediction using the Rabani et al. UTR-seq benchmark.

| Model | Pearson $r$ | Spearman $\rho$ | $R^2$ | MAE |
| --- | --- | --- | --- | --- |
| EukaUTR-S2 | **0.688 ± 0.007** | **0.681 ± 0.007** | **0.449 ± 0.032** | **0.156 ± 0.007** |
| EukaUTR-S2-LoRA | *0.684 ± 0.006* | *0.676 ± 0.008* | *0.445 ± 0.037* | 0.158 ± 0.009 |
| EukaUTR-S1 | *0.684 ± 0.009* | *0.676 ± 0.008* | 0.442 ± 0.035 | 0.158 ± 0.008 |
| 3UTRBERT | 0.677 ± 0.008 | 0.670 ± 0.008 | 0.407 ± 0.048 | 0.161 ± 0.007 |
| mRNABERT | 0.674 ± 0.009 | 0.669 ± 0.008 | 0.437 ± 0.014 | *0.157 ± 0.003* |
| GEMORNA-UTR | 0.594 ± 0.006 | 0.579 ± 0.007 | 0.327 ± 0.019 | 0.175 ± 0.004 |

Reporter half-life was derived for sequences with valid positive degradation-rate estimates as $t_{1/2} = \ln(2)/\beta$ . Values are mean ± s.d. across ten held-out test folds. Bold and italic type indicate the best and second-best values for each metric, respectively; ties share formatting. Pearson $r$ , Spearman $\rho$ and $R^2$ are higher-is-better metrics, whereas MAE is lower-is-better.

**Supplementary Table 13.**
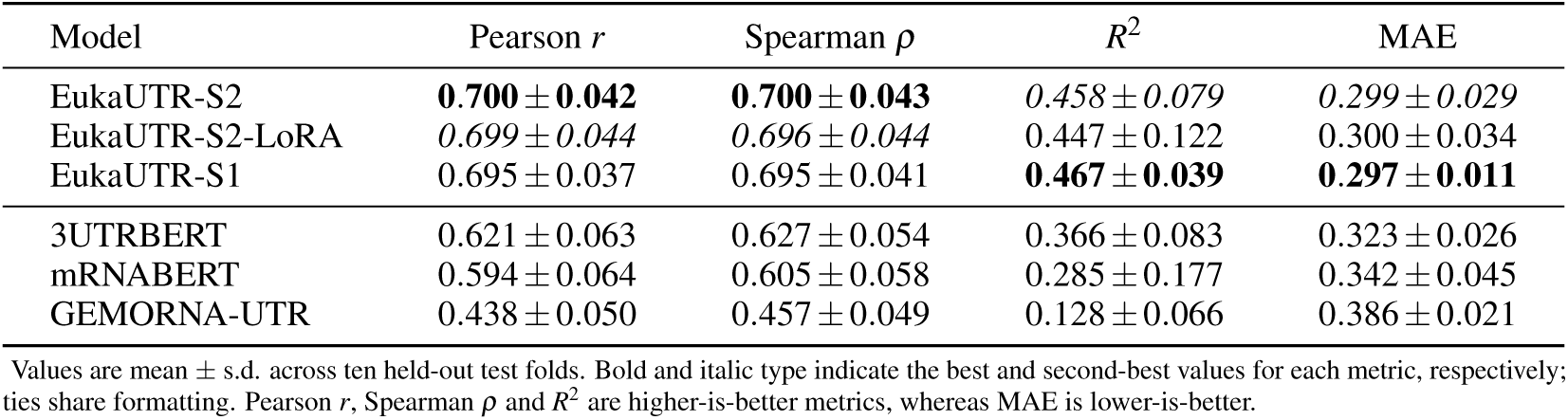
Performance on human reporter mRNA half-life prediction using the Zhao et al. fast-UTR benchmark.

| Model | Pearson $r$ | Spearman $\rho$ | $R^2$ | MAE |
| --- | --- | --- | --- | --- |
| EukaUTR-S2 | **0.700 ± 0.042** | **0.700 ± 0.043** | *0.458 ± 0.079* | *0.299 ± 0.029* |
| EukaUTR-S2-LoRA | *0.699 ± 0.044* | *0.696 ± 0.044* | 0.447 ± 0.122 | 0.300 ± 0.034 |
| EukaUTR-S1 | 0.695 ± 0.037 | 0.695 ± 0.041 | **0.467 ± 0.039** | **0.297 ± 0.011** |
| 3UTRBERT | 0.621 ± 0.063 | 0.627 ± 0.054 | 0.366 ± 0.083 | 0.323 ± 0.026 |
| mRNABERT | 0.594 ± 0.064 | 0.605 ± 0.058 | 0.285 ± 0.177 | 0.342 ± 0.045 |
| GEMORNA-UTR | 0.438 ± 0.050 | 0.457 ± 0.049 | 0.128 ± 0.066 | 0.386 ± 0.021 |

Values are mean ± s.d. across ten held-out test folds. Bold and italic type indicate the best and second-best values for each metric, respectively; ties share formatting. Pearson $r$ , Spearman $\rho$ and $R^2$ are higher-is-better metrics, whereas MAE is lower-is-better.

**Supplementary Table 14.**
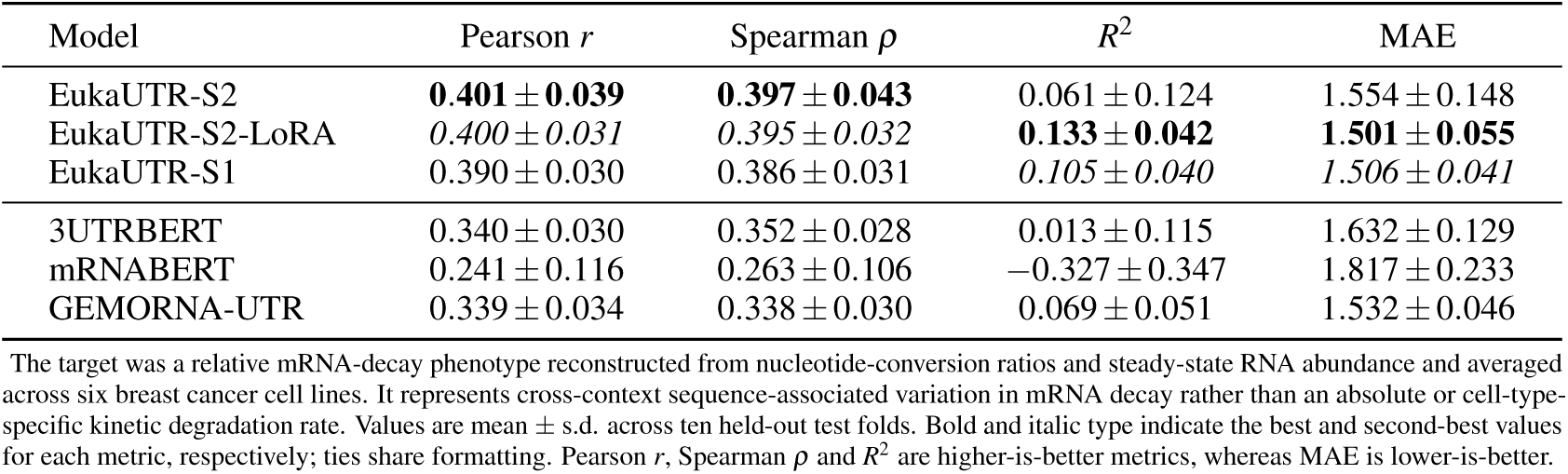
Performance on the cross-context relative mRNA-decay benchmark reconstructed from breast cancer SLAM-seq data.

**Supplementary Table 15.**
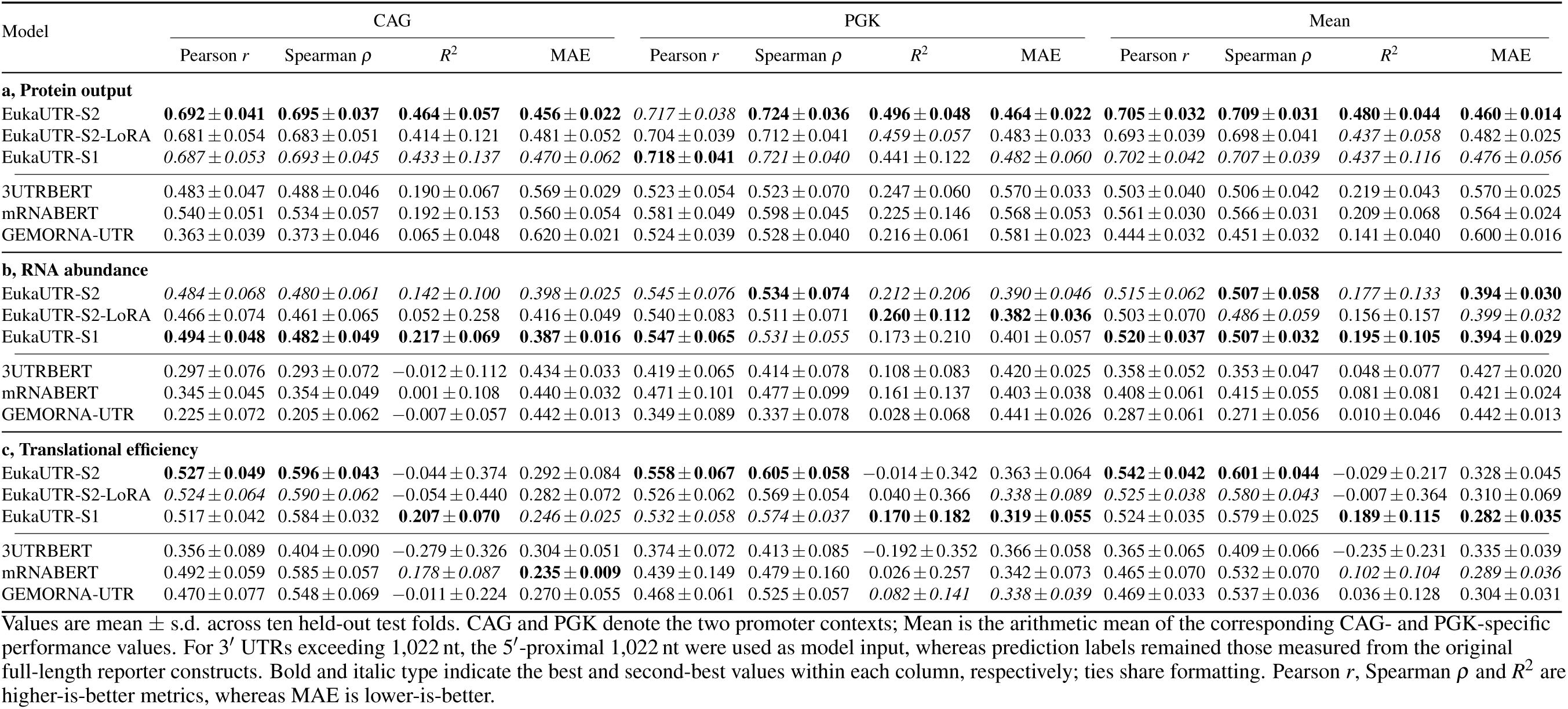
Performance on full-length human 3′ UTR regulatory-output prediction across CAG and PGK promoter contexts.

**Supplementary Table 16.** Performance on the viral MPRA RNA-abundance and mean-ribosome-load benchmarks.

| Model | Pearson $r$ | Spearman $\rho$ | $R^2$ | MAE |
| --- | --- | --- | --- | --- |
| **RNA abundance** |  |  |  |  |
| EukaUTR-S2 | *0.472 ± 0.039* | *0.385 ± 0.052* | *0.167 ± 0.094* | 0.210 ± 0.023 |
| EukaUTR-S2-LoRA | 0.461 ± 0.046 | **0.386 ± 0.055** | 0.165 ± 0.065 | *0.205 ± 0.015* |
| EukaUTR-S1 | **0.481 ± 0.050** | 0.382 ± 0.053 | **0.171 ± 0.088** | 0.208 ± 0.018 |
| 3UTRBERT | 0.426 ± 0.068 | 0.364 ± 0.066 | 0.143 ± 0.054 | **0.204 ± 0.020** |
| mRNABERT | 0.359 ± 0.072 | 0.325 ± 0.056 | 0.106 ± 0.043 | 0.210 ± 0.017 |
| GEMORNA-UTR | 0.333 ± 0.068 | 0.291 ± 0.067 | 0.100 ± 0.050 | 0.211 ± 0.019 |
| **Mean ribosome load (MRL)** |  |  |  |  |
| EukaUTR-S2 | 0.265 ± 0.117 | 0.178 ± 0.023 | *−0.004 ± 0.118* | 0.200 ± 0.018 |
| EukaUTR-S2-LoRA | *0.276 ± 0.109* | *0.180 ± 0.027* | *−0.009 ± 0.148* | *0.199 ± 0.020* |
| EukaUTR-S1 | **0.278 ± 0.100** | **0.183 ± 0.025** | **0.008 ± 0.121** | **0.198 ± 0.021** |
| 3UTRBERT | 0.267 ± 0.087 | 0.164 ± 0.025 | *−0.082 ± 0.116* | 0.212 ± 0.023 |
| mRNABERT | 0.184 ± 0.064 | 0.155 ± 0.024 | *−0.031 ± 0.074* | 0.199 ± 0.027 |
| GEMORNA-UTR | 0.203 ± 0.084 | 0.132 ± 0.027 | *−0.080 ± 0.201* | 0.205 ± 0.020 |

RNA abundance and mean ribosome load (MRL) were evaluated as separate quantitative outcomes for 130-nt viral genomic tiles inserted into the 3' UTR of a common reporter. Values are mean ± s.d. across ten held-out test folds grouped by viral genome accession, such that tiles derived from the same accession did not cross training and test partitions. Bold and italic type indicate the best and second-best values for each metric, respectively; ties share formatting. Pearson $r$ , Spearman $\rho$ and $R^2$ are higher-is-better metrics, whereas MAE is lower-is-better. MRL, mean ribosome load.

